# Available time-series of terrestrial animals provide low reliability in population trend estimates

**DOI:** 10.64898/2026.10.14.759066

**Authors:** Julia G. de Aledo, Ignasi Bartomeus, François Duchenne

## Abstract

Monitoring biodiversity through time is critical for understanding the impacts of global change, yet the reliability of available time-series for estimating population trends remains poorly studied. We simulated population dynamics to study how monitoring duration, inter-annual variability, and temporal autocorrelation influence trend reliability. We then inferred the expected reliability of 16,450 empirical time-series of terrestrial animals based on those simulations. Longer, less variable, and negatively autocorrelated time-series produced more reliable trend estimates. Time-series shorter than 20 years led to large magnitude errors and low power in estimating population trends, despite generally low sign uncertainty. Overall, 80% of the empirical time-series were too short and/or noisy to identify meaningful temporal trends, yet reliability varies among taxa. Our results show important limitations in the available biodiversity data, stressing the need to sustain long-term monitoring to better estimate population trends, a top priority in the current biodiversity crisis.

## Introduction

Reliable estimates of animal population change are fundamental for understanding biodiversity trends and evaluating conservation policies. During the last decades, scientific, societal and political concerns regarding global biodiversity loss have increased substantially. Biodiversity has been reshaped by human-induced rapid environmental change (Jaureguiberry et al., 2022; Sala et al., 2000), which led to declines in species abundance (Ceballos et al., 2017) and local species richness (Murphy & Romanuk, 2014), as well as changes in community composition (Dornelas et al., 2014). However, quantifying the magnitude and direction of biodiversity change remains a challenge. While global extinction rates continue to rise (Ceballos & Ehrlich, 2023; Dirzo & Raven, 2003; Pimm et al., 2014), synthesis of local diversity often provides contrasting results (Cardinale et al., 2018), complicating both ecological understanding and conservation planning (Valdez et al., 2023).

In 2022, the Kunming–Montreal Global Biodiversity Framework fixed ambitious goals, such us halting extinctions and maintaining healthy populations by 2030 (Convention on Biological Diversity, 2022). Evaluating the success of conservation policies within such short time scales requires trustable indicators of population change to detect whether observed declines are slowing down or being reversed. Different indicators have emerged to monitor changes in species abundance across time, such as the Living Planet Index (Collen et al., 2009), the bird farmland index (PECBMS, 2025) or indicators using butterfly monitoring schemes (Edwards et al., 2025). These indicators and data bases have now accumulated decades of data, but they remain restricted to specific groups such as birds, butterflies or mammals. Time-series on other taxa consist on heterogeneous collections of available data, such as the ones on BioTIME (Dornelas et al., 2018, 2025), which often encompass data collected with a wide range of sampling methodologies and durations. These aggregations have been precious for assessing biodiversity changes across the last decades (Chaikin et al., 2024; PECBMS, 2025). However, it remains unclear if these data can allow us to estimate reliable trends, particularly over the short timescales at which conservation policies are evaluated.

Reliability analyses are a common practice in ecology, especially in the design of experiments (Steidl & Thomas, 2001). However, in observational studies that do not control the environment and use complex statistics to separate confounding effects, reliability analyses become much more complex to design (Johnson et al., 2015). Yet, these analyses have been widely overlooked over the last decades –but see Leung & Gonzalez, 2024; Yang et al., 2022–. Consequently, weak increases or declines can be ignored when statistical power is low (type II error; White, 2019), whereas considering significant results only is likely to overestimate the magnitude of population change (type M error; Gelman & Carlin, 2014). Reliability analyses can be easily estimated by assessing few of the time-series characteristics.

First, the duration of the time-series is one of the main indicators of reliability. The longer the time-series, the higher the reliability of the estimated temporal trend (Gerrodette, 1987; White, 2019). Some studies have pointed out the over-representation of recent and short time-series in the available data (Duchenne et al., 2022; Loreau et al., 2022; Mihoub et al., 2017). This means that recent monitoring programs would tend to have low reliability, not being able to detect weak declines or detecting trends that are actually non-existent; questioning the reliability of these time-series to detect population changes.

Second, the inter-annual variability around the average trend, sometimes referred to as noise, stochasticity or population variability, also affects the reliability of the estimated temporal trend (Wilson et al., 2011). The higher the inter-annual variability, the lower the reliability (Gerrodette, 1987), hiding existing temporal trends or interpreting transient increases or decreases as signs of true long-term trends (Connors et al., 2014). Since inter-annual variability depends on taxa (Montgomery et al., 2026) and environmental conditions (among others), the statistical power to detect changes varies across taxonomic groups, space, and time (Dove et al., 2023; Leung & Gonzalez, 2024; White, 2019).

Third, the effect of inter-annual variability on the reliability of estimated temporal trends can be further exaggerated when it is autocorrelated in time (Connors et al., 2014). Autocorrelated variability can arise from the abiotic environment (Engen et al., 2013; Nicolau et al., 2022), species interactions (Dallas & Kramer, 2022; Hansen et al., 1999), age-structured dynamics (Vindenes & Engen, 2017), or the intrinsic dependence of population size on its previous state, a fundamental property of density-regulated population dynamics (Dennis et al., 2016), affecting survival rates and abundance. The strength and direction of temporal autocorrelation may also vary among taxa, reflecting differences in population dynamics and responses to environmental fluctuations.

In order to address our current ability to estimate reliable population trends, we first used simulations to evaluate the relative contribution of time-series characteristics – duration, inter-annual variability and temporal autocorrelation– in determining the reliability of estimated population trends. Then, we combined our simulation results with the recently released compilation of empirical time-series data from BioTIME 2.0 database (Dornelas et al., 2025) to extract the expected reliability in estimated population trends offered by time-series for each taxonomic group (amphibia, arachnids, birds, insects, mammals and squamates). Our results show that available time-series of terrestrial animals offer low reliability in estimating population trends, questioning our ability to monitor and predict biodiversity changes.

## Methods

We first simulated population abundance time-series spanning the ranges of duration, inter-annual variability and temporal autocorrelation observed in the BioTIME data for terrestrial animals. We compared the estimated parameters obtained from modelling with the true values used to generate them to quantify five reliability metrics (estimation error, Type M error, Type S error, statistical power, false-positive rate). We then extracted the time-series characteristics of the empirical data to assess their expected reliability (Fig. 1).

**Fig. 1.**
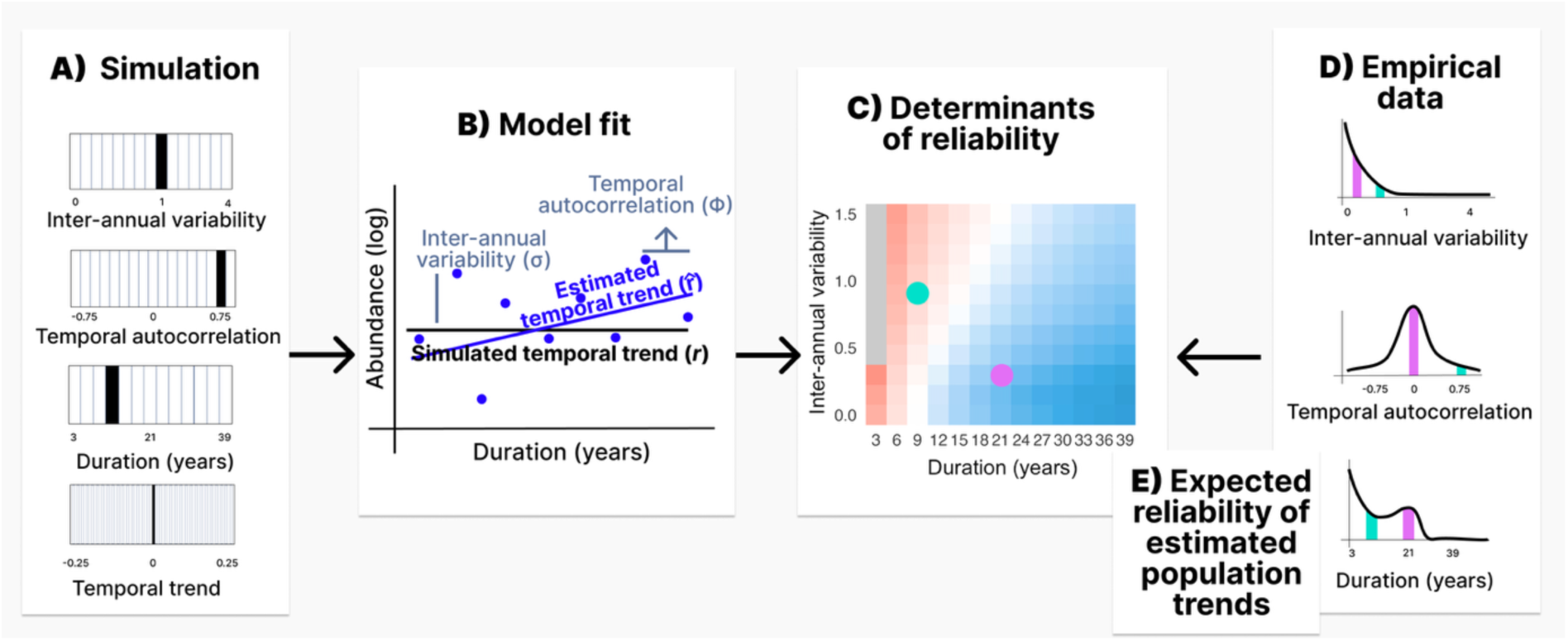
Schematic representation of our study workflow. **(A)** We first selected a range of time-series characteristics based on empirical data to build the simulated data. Empty lines show the different values used to generate simulations and the thick lines show the parameter value used for the example illustrated in (B). **(B)** For each combination, we simulated yearly abundances (blue points) drawn from log-normal Poisson distribution. Abundance was changing on average across years with a true temporal trend (r, black horizontal line), exhibited a certain inter-annual variability (σ) and could be autocorrelated with a certain strength (Φ). We ran GLMMs on simulated data to estimate temporal trends (*r^*) (blue line) and compare them with the real trend. **(C)** The reliability of the temporal trends was assessed by different reliability metrics as a function of all time-series characteristics, colour gradient represent the estimation error (from dark blue 0%.year^-1^ to dark red 25%-year^-1^error) and **(D)** using BioTIME 2.0 filtered by terrestrial animals, we obtained their time-series characteristics with GLMMs for each species models (pink and turquoise bars represent the characteristics of two random empirical time-series) and **(E)** we inferred the expected reliability in population trend estimation by locating them (e.g. pink and turquoise points) according to their time-series characteristics and extracting their reliability metrics from the reliability surface (C).

### Time-series simulation

Our simulation process allows for an independent inclusion of a temporal trend, its inter-annual variability and temporal autocorrelation while generating count data. To implement these simulations, first, we model the deterministic variation of population size across time as a geometric series, which allowed us to simulate stable, declining or increasing populations. Population size (µ*_t_*), at year *t* is defined as follows:

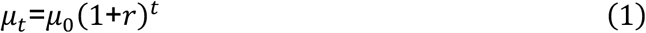

Where µ_0_ is the initial average number of individuals and *r* is the growth rate of the population. Then, we simulated a stochastic variation around that process to simulate yearly individual counts with a given inter-annual variability. Note that here we do not focus on the sources of the inter-annual variability, population dynamics and/or to sampling error, but on the consequences of such variability, regardless its determinants. To simulate observed individual counts, for each year *t*, a number of individuals (*Y_t_*) is drawn from a Poisson-lognormal distribution, for which the mean *λ_t_* is drawn from a log-normal distribution with a mean *log*(*μ_t_*):

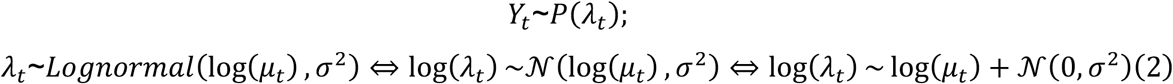

where *σ*^2^ is the variance of the normally distributed deviations on the logarithmic scale. This way, our simulation can account for the inter-annual variability through the overdispersion of the Poisson distribution. Inter-annual variability was introduced through stochastic variation in the expected annual abundance, generating overdispersion in observed counts relative to a Poisson distribution. To implement temporal autocorrelation in the inter-annual variability, we used a multivariate Normal distribution to draw simultaneously all the *λ_t_*, stored in a vector λ⃗*_t_*:

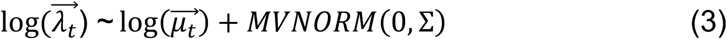

where *log*(μ⃗) is the vector of the logarithm of the yearly average abundances (*μ_t_*) and Σ is the variance-covariance matrix between yearly abundance values, built according to an autoregressive process of order one (AR1). The covariance between year *t* and year *k* being defined as follows:

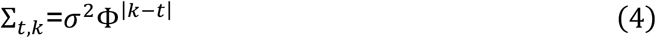

Where *σ*^2^ is the variance across years in the values of *λ_t_* and thus measures the inter-annual variability, while Φ is the temporal autocorrelation in that inter-annual variability between two consecutive years. For interpretability, we used the standard deviation *σ* = √*σ*^2^, which can be seen as an overdispersion parameter. An inter-annual variability of *σ* = 0 means we used a Poisson distribution. The higher *σ*, the higher the overdispersion relative to a Poisson distribution. To give an interpretable scale to the inter-annual variability, *σ* can be linked to the magnitude of the inter-annual variability (*τ*) relative to the average abundance: *τ* = *e*^1.96σ^. An inter-annual variability of *σ* = 0.1 means an inter-annual variability up to *τ* ≈ 1.22 times the deterministic average abundance (*μ_t_*). If the average population size is *μ_t_* = 100, this means that the inter-annual variability would tend to produce values that have 95% of chances to included in O<u>^1^</u> 100,100*τ*P = [82,122].

To perform simulations, we selected ranges of time-series characteristics that covered most of the available time-series from BioTIME 2.0 database (cf. below). We simulated population dynamics for 14 values of duration (from 3 to 39 years each 3 years, plus an additional value of 57 years), 15 values of inter-annual variability (*σ* ranging from 0 to 4) and 13 values of temporal autocorrelation (Φ ranging from −0.9 to 0.9 in increments of 0.15). This led to 2730 combinations of time-series characteristics. For each of these, we simulated for 51 different temporal trends (ranging from −0.25 to 0.25 in increments of 0.01 (i.e. 1%.year^-1^), using 100 stochastic replicates with a different initial abundance (sampled between 1 and 90), which led to 5100 simulated time-series per combination of time-series characteristics, leading to a total of 13,923,000 simulated time-series.

### Assessing reliability of temporal trends estimated from simulated time-series

To assess the reliability that simulated time-series offer in the inference of the temporal trend in abundance, we compared the temporal trend estimated with generalized linear mixed-effect models (GLMM) with the true population trend (*r*) from the simulated time-series. We fitted one GLMM with a Poisson error-structure, and a log link function per time-series, using a structure that closely fitted the simulation process, with a linear effect of the years and a year random effect to account for the inter-annual variability and its temporal autocorrelation:

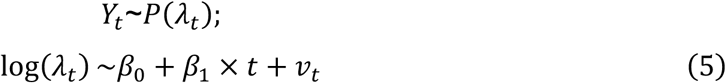

where the year abundance (*λ_t_*) was modelled as a function of an intercept (*β*_0_) and a linear effect of the year (*β*_1_), as well as a random effect of the year (*v_t_*). This random effect is normally distributed, with a variance-covariance matrix (Σ) structured according to an AR1 process, to capture potential temporal autocorrelation:

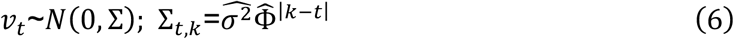

where 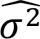 is the estimated variance of inter-annual variability and *Φ*^Z^ is the estimated strength of the temporal autocorrelation. Note that there is a direct correspondence between the linear effect of the year from the GLMM and the population growth rate (Duchenne et al., 2022): *r*^ = *e^β^*^^1^ − 1, where *r*^ is the estimated growth rate that can be directly compared with *r*.

Models that failed to meet standard convergence and diagnostic criteria were excluded, retaining 12,616,022 models. Non-convergence was concentrated in three-year time-series, for which some parameter combinations lost more than 4,800 of the 5,100 simulated models per combination of time-series characteristics (Fig. S1). To ensure sufficient replication, combinations of time-series characteristics retaining fewer than 20% successfully fitted models (n = 1020) were eliminated from subsequent analyses. We excluded 72 combinations of time-series characteristics out of 2730, all of which corresponded to very short durations (3 years), explaining why models did not converge.

To quantify the reliability of the statistical inference, we used five different metrics: (i) the error in the estimated temporal trend, defined as the absolute difference between the estimated and the true population temporal trend, *E[|ř − r|]* (estimation error from now onwards); (ii) Type M error, which measures the degree to which statistically significant estimates exaggerate or underestimate the true trend 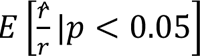; where *p* is the p-value associated with *r^* (i.e. *β*^c^_1_); (iii) Type S error: is the probability that the model estimates a trend in the opposite direction of the true trend, *P*(*sign*(*r^*) ≠ *sign*(*r*)|*p* < 0.05) (Gelman & Carlin, 2014); (iv) the statistical power, defined as the probability that the model correctly detects a given temporal trend, *P*(*p* < 0.05 ∧ r^ < 0 ∣ *r* = −0.02) where *r*_0_ in this study is −0.02 (−2%.year⁻¹), and (v) the false-positive rate (Type I error), defined as the probability of detecting a statistically significant trend when the true trend is zero, *P*(*p* < 0.05 ∣ *r* = 0). We defined thresholds of acceptable reliability for each metric used: for error, we used a threshold of 2%.year⁻¹ to limit the average deviation from the real temporal trend below 2%.year⁻¹. A deviation from 2%.year⁻¹ reduces by half a population in 35 years, and thus such deviation is not negligible. We defined acceptable statistical power when a decline of 2%.year^-1^ was detected in 80% of the cases, following conventional standards (Cohen, 1988). False-positive rate was acceptable under 5%, the accepted significance level (α = 0.05). Type S error is expected to be very low and thus defined as acceptable at 1% error maximum. Type M error was considered acceptable below 1.2 (i.e. <20% exaggeration), following recommendations (Gelman & Carlin, 2014). We also evaluated the potential bias in the estimation of temporal autocorrelation (*Φ*^^^ − *Փ*), inter-annual variability (*σ*k − *σ*), and temporal trend (*r*^-*r*) across all combinations of simulated time-series characteristics (Fig. S2). We also calculated the Pearson correlations among the five reliability metrics and the three time-series characteristics to assess their pairwise associations (Fig. S3).

### Assessing reliability of statistical inference offered by empirical time-series

#### Empirical time-series

We used population time-series from BioTIME 2.0 (Dornelas et al., 2025) because it has been one of the most used databases to study population changes (Chaikin et al., 2024; Daskalova et al., 2020; Justin Nowakowski et al., 2023). We target terrestrial species from the groups Amphibia, Arachnida, Aves, Insecta, Mammalia, and Squamata. Taxonomic identities were resolved using GBIF backbone taxonomy (Chamberlain et al., 2025). Since sampling resolution differed among studies (Dornelas et al., 2025), we harmonized spatial units prior to analysis. Population counts originating from the same study were aggregated into spatial clusters using hierarchical clustering on geographic distances. Since spatial grain must serve as a proxy for taxa mobility (Valdez et al., 2023), we used a 5 × 5 km resolution for birds and mammals, 2 × 2 km for squamates and amphibians, and a 1 × 1 km resolution for insects and arachnids, reflecting typical insect sampling protocols (Gaume & Desquilbet, 2024).

Over the 22,577 filtered time-series, we retained only consecutive time-series of abundance data that were at species level, with a minimum of three years and with at least one positive abundance estimate. After filtering, the dataset comprised 16,450 time-series, regarding 2,739 species, spread across the six groups: amphibia (109 species), arachnids (96), birds (612), insects (1660), mammals (162), and squamates (100). Retained time-series were mostly spread in North America, Europe and Japan (Fig. S4). Since BioTIME 2.0 stored presences only, we imputed a zero when a species was not detected in a given year, study and location while at least one other species was detected in the same year, study and location, following the established protocol for reconstructing pseudo-absences (Dornelas et al., 2025).

#### Assessing expected reliability

To assess the expected reliability offered by empirical time-series, we first needed to assess the empirical time-series characteristics. Duration was defined by the number of years the monitoring was held on (Fig. S5); inter-annual variability and temporal autocorrelation were estimated using the same model that we applied to the simulated time-series, described by equations (5) and (6). Finally, we matched the extracted time-series characteristics to the closest corresponding values we had in our simulations to obtain associated values of the five reliability metrics. 454 time-series (2.7%) fell within the parameter space for which reliability metrics have not been estimated due to low sample size (grey cells in Fig. 2), and were removed from the analysis, retaining 15,996 time-series.

**Fig. 2.**
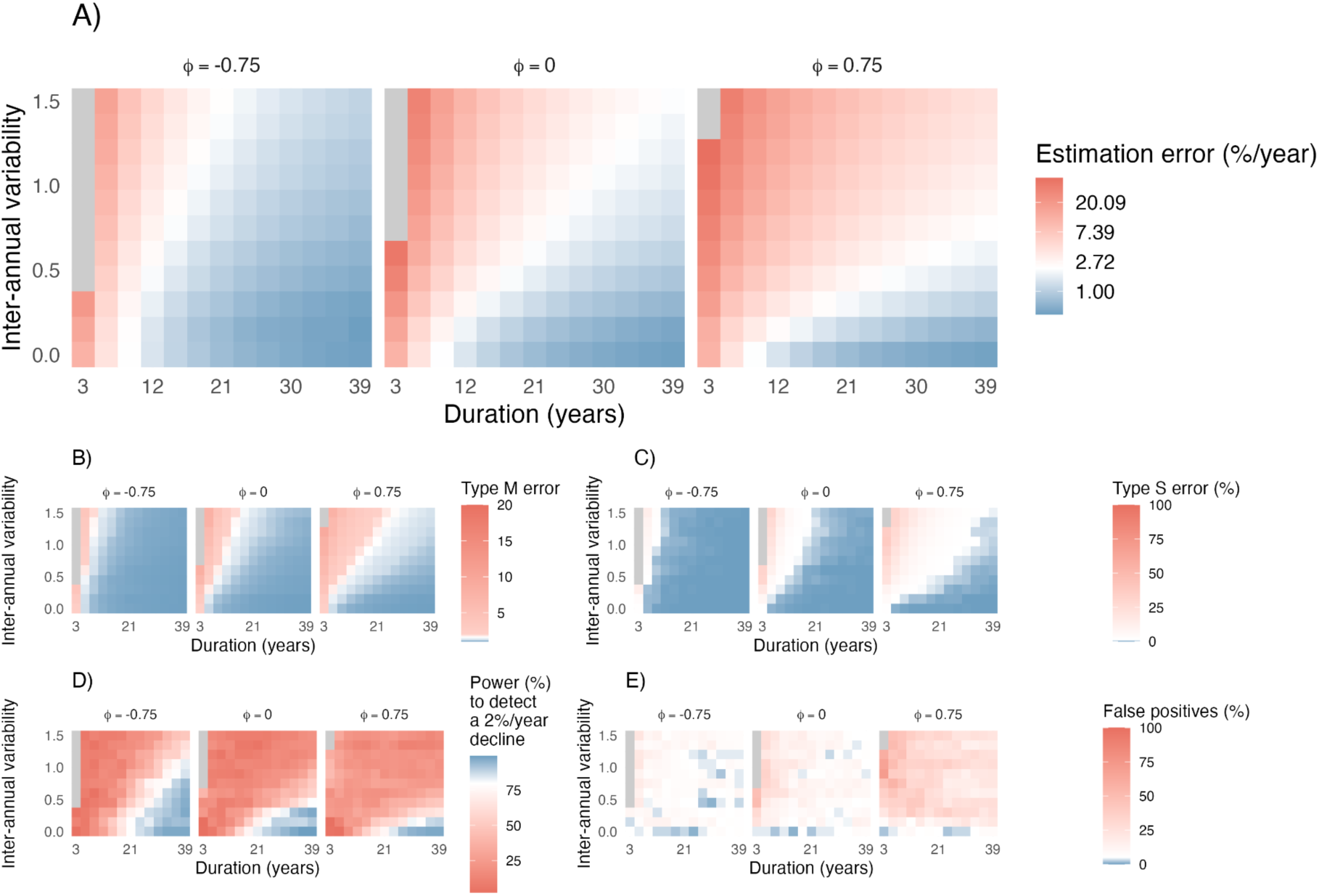
Reliability of temporal trend estimation varies across time-series characteristics. Heatmaps show five reliability metrics associated with temporal trend estimation: **(A)** mean estimation error, **(B)** Type M error (magnitude exaggeration conditional on significance), **(C)** type S error (probability of inferring the incorrect trend direction), **(D)** statistical power to detect a decline of 2%.year^-1^, **(E)** false-positive rate. Metrics are shown as a function of time-series duration, inter-annual variability (σ), and temporal autocorrelation (Φ). For clarity, heatmaps are presented for three representative autocorrelation values (Φ = −0.75, 0, and 0.75), corresponding to strongly negative, neutral, and strongly positive temporal autocorrelation. Cells representing less than 20% of converging models are shown in grey. For each color scale, white has been centered on the threshold defining acceptable reliability (estimation error = 2%.year⁻¹, Type M = 1.2, Type S = 1%, power = 80%, and false-positive rate = 5%).

## Results

### The effect of time-series characteristics on the reliability of temporal trends

Our results show that the reliability of estimated temporal trends varied strongly with duration, inter-annual variability and temporal autocorrelation (Fig. 2). Most three-year time-series were too short to infer reliable trends, mostly under negative autocorrelation (Fig. 2 & S1) and were removed from the analysis. Although duration appears to be the main driver of reliability (Fig. S2), analysing long time-series did not guarantee to infer reliable population trends. Reliability strongly depended on inter-annual variability and its temporal autocorrelation, meaning that even 40 years long time-series could offer low reliability when inter-annual variability and autocorrelation were high.

Estimation error, type M, type S errors were positively correlated among them and highly related to the time-series’ duration (Fig. S3). Power was driven by duration and inter-annual variability, while the rate of false-positives was mostly determined by the temporal autocorrelation of inter-annual variability (Fig. 2A-D & S3). In general terms, type S tended to remain low, indicating that it is unlikely to infer a significant trend in the opposite direction of what it is in reality. However, < 20 years and variable time-series led to very large errors in the estimation of the trend, exceeding 20%.year^-1^ error (Fig. 2A), meaning that this rate of decline could easily be mistaken for a population without any trend. Moreover, these time-series often led to an inflated type M error, suggesting an overestimation of the magnitude of population trends up to twelve times (Fig. 2B). Yet, when increasing duration above 20 years, coupled with a moderate variability, time-series could reach satisfying levels of estimation error, type M and type S errors (Fig. 2C). However, reaching a satisfying statistical power was the hardest condition to satisfy: it required long time-series with very low inter-annual variability, especially if inter-annual variability was positively autocorrelated through time (Fig. 2D & S5).

Contrary to other reliability metrics, false-positive rates were primarily influenced by temporal autocorrelation, with a lower influence of the duration or the strength of the inter-annual variability. Since negative temporal autocorrelation creates a symmetric and homogeneous abundance variation around the trend –where favourable and unfavourable years tend to alternate–, it helped inference and increased reliability (Fig. 2E). In contrast, positive autocorrelation tended to create heterogeneous variations of abundance around the average trend, with temporal windows with positive abundance deviation, relative to the mean, and other windows with negative deviations. This decreased the reliability of the estimated trend and was likely to create false-positives, exceeding 25% under many conditions (Fig. 2E).

### Characterizing empirical time-series

The empirical time-series extracted from BioTIME 2.0 showed considerable heterogeneity in duration and temporal structure among taxonomic groups (Fig. 3). Time-series duration varied widely: amphibians, squamates and arachnids exhibited low mean durations of approximately 5 to 6.5 years, mammals around 9.5 years, insects 14.5 years, and birds, as expected, showed the longest duration with 22 years. For inter-annual variability, amphibians and squamates showed low variability (σ^ = 0.300, σ^ = 0.297 respectively), with variation that reached on average 1.8 times the average abundance. Arachnids and birds displayed intermediate variability (σ^ = 0.422, σ^ = 0.463 respectively), with variation in abundance being around 2.3 times the average abundance, while mammals showed slightly higher values (*σ*k = 0.686). Finally, insects exhibited the highest mean inter-annual variability (*σ*k = 0.794), with variation in abundance reaching 4.7 times the average abundance, reflecting substantial year-to-year fluctuations. Mean temporal autocorrelation was generally close to 0 for all taxa. Arachnids, mammals, and squamates showed slightly negative mean autocorrelation (Φ^Z^ = −0.25). By contrast, birds tended to show a positive autocorrelation on average (Φ^Z^ = 0.08), whereas insects had values centred on zero (Φ^Z^ = −0.05).

**Fig. 3.**
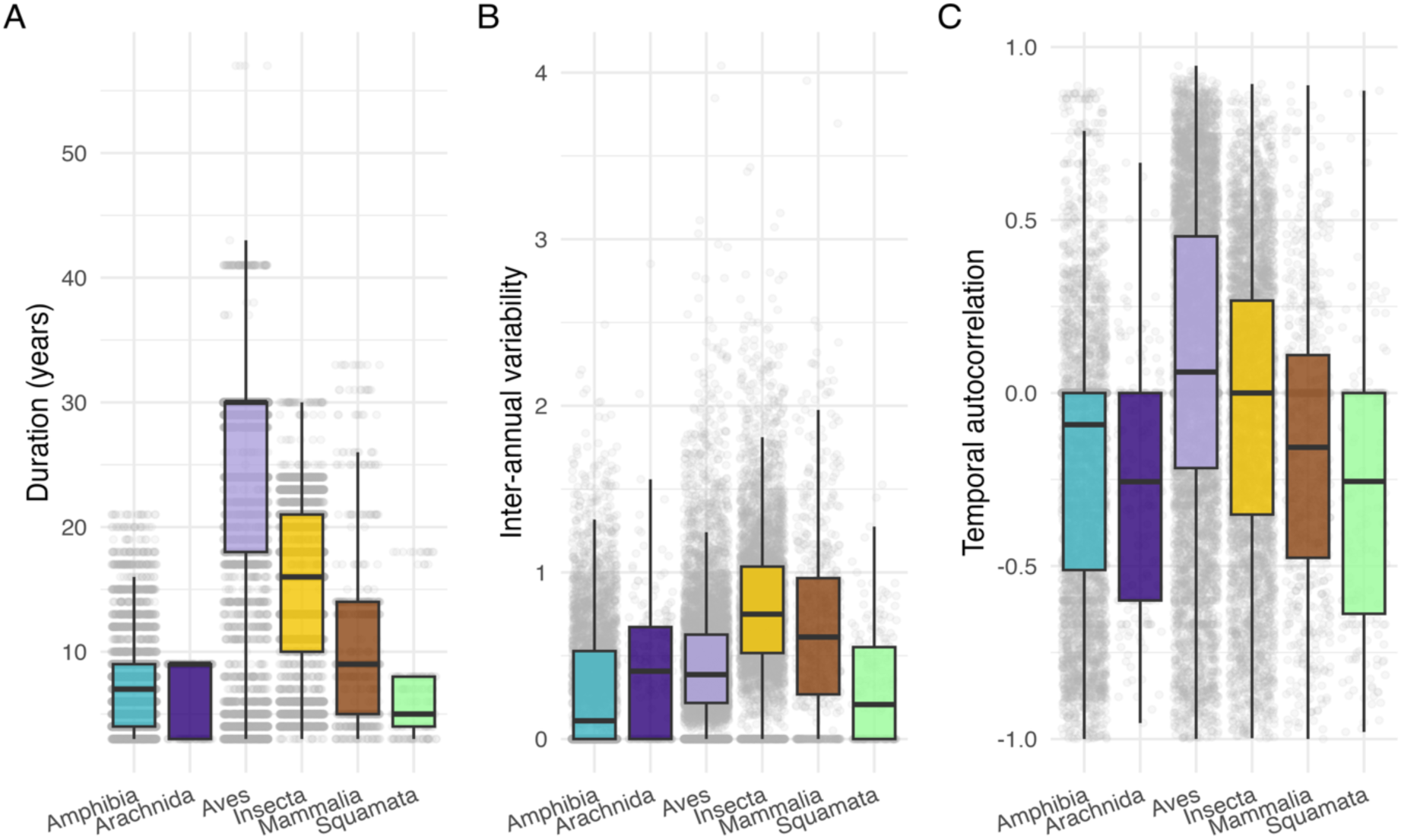
Characteristics of empirical time-series available in BioTIME for terrestrial animals. **(A)** duration of time-series (in years), **(B)** inter-annual variability, **(C)** temporal autocorrelation, as a function of taxonomic groups. Points represent values for each time-series, while boxplots represent the distribution of these values.

### Assessing statistical reliability offered by empirical time-series

Reliability differed among taxa (Fig. 4): bird time-series consistently offered higher reliability by exhibiting the lowest expected estimation errors (Fig. S7), the lowest magnitude exaggeration, and the highest statistical power. In contrast, time-series from amphibians, mammals, squamates, and especially insects were generally associated with time-series characteristics offering less reliable population trends. Consistently with previous findings (Cortés-Guzmán et al., 2026; Yang et al., 2022), type M error was substantial for most taxa (Fig. 4B), indicating that the magnitude of population change is likely to be exaggerated if interpreting only significant results. In addition, statistical power was also strongly limited (Fig. 4D): only a small fraction of time-series was expected to achieve sufficient power to detect a decline of 2%.year^-1^, suggesting that gradual population declines frequently remain undetected. This is consistent with previous results (Wauchope et al., 2019; White, 2019): they misestimate gradual population declines, producing either non-significant or extreme population trends. In addition, the expected false-positive rate exceeded conventional expectations of 5% for most of the time-series for all taxa, indicating an elevated risk of inferring spurious trends. Together, these results suggest that although the direction of temporal change can often be inferred correctly (low type S error), the magnitude of change and the ability to reliably detect gradual declines, if they occur, remain major challenges for most empirical time-series.

**Fig. 4.**
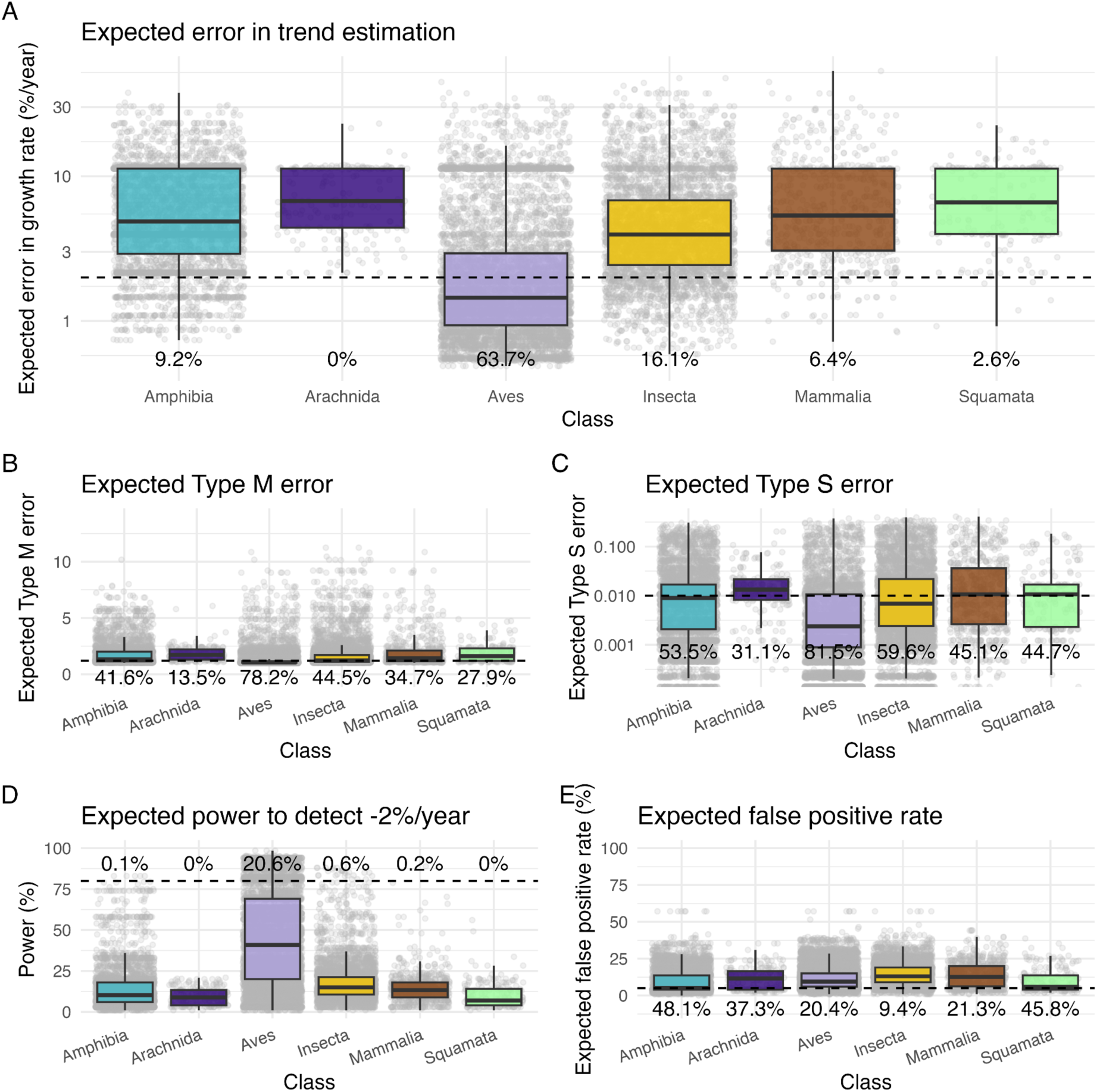
Expected reliability of temporal trends estimated from empirical time-series. Each 15,996 time-series from BioTIME 2.0 was matched to the closest simulation scenario based on its duration, autocorrelation, and inter-annual variability. Expected **(A)** error in trend estimation, **(B)** type M error, **(C)** type S error, **(D)** power to detect a 2%.year^-1^ decline and **(E)** false-positive rate of the data available per taxa (Amphibia, Arachnida, Birds, Insects, Mammals and Squamata). Points represent individual time-series, while boxplots represent the distribution of these values. Text shows the percentage of time-series within the expected threshold (error < 2% year⁻¹, Type M < 1.2, Type S < 1%, power > 80%, and false-positive rate < 5%).

## Discussion

Our results show that ecological time-series are often too short and noisy for inferring reliable population trends from the time-series we currently have. Thus, our work suggests that before interpreting estimated population trends, researchers should carefully evaluate the characteristics of the time-series they study, as they might affect the robustness of the conclusions. This also stresses the need for a stronger investment in long-term biodiversity monitoring, a key strategy for improving the reliability of population trend estimates that can be complemented by other approaches.

### Time-series characteristics effect on trend reliability

Our simulations confirm the central role of duration, inter-annual variability, and temporal autocorrelation on the reliability of ecological time-series. Previous studies have focused on how duration affects statistical power (White, 2019) or the error in the estimation of the temporal trends (Wauchope et al., 2019). Here we completed those studies by extending the simulations to different levels temporal autocorrelation and by evaluating simultaneously multiple facets of reliability. Our results highlight that reliability metrics can be grouped in three facets of reliability according to how they covary. First, estimation error, type M and type S errors were highly correlated, relating to the precision of the estimation primarily driven by duration. Second, statistical power on its own represents the second facet, related to our ability to detect changes, driven by duration and inter-annual variability with equal importance. Finally, the third was represented by false-positives rate, driven mainly by temporal autocorrelation.

Duration emerges as the one of the primary characteristics affecting reliability. For example, statistical power increased with monitoring length, with 19.5 years as an average minimum duration to reach the 80% power and with less than six years duration being useless to detect minor or moderate declines (Fig S5), both findings being highly consistent with previous studies (Wauchope et al., 2019; White, 2019). Moreover, the negative effect of inter-annual variability on reliability also depended on the duration: reliability dropped more sharply with inter-annual variability for short time-series than for long time-series. Increasing monitoring duration is thus an important factor to improve estimated temporal trends, especially for taxa that exhibit high inter-annual variability in population size.

Temporal autocorrelation represented a third, and often disregarded, source of uncertainty (Daskalova et al., 2021; Johnson et al., 2024). We found that negative autocorrelation made trends easier to detect because it creates a homogeneous, symmetric noise around the average abundance, thus helping the statistical inference. In contrast, positive autocorrelation decreased the reliability of time-series because it creates time dependencies that increase the risk of detecting trends under stable conditions. Autocorrelated low or high values, particularly at the beginning or end of the time-series, can exert disproportionate leverage (Duchenne et al., 2022).

### Reliability offered by empirical time-series

Previous studies have focused on how the duration of time-series affects statistical power or estimation error when estimating temporal trends for birds (Wauchope et al., 2019), fish and mammals (White, 2019), or river invertebrates (Cortés-Guzmán et al., 2026). Here, we complete these studies by performing an exhaustive analysis of the reliability offered by terrestrial animals’ available time and considering five reliability metrics. Our results highlight that most of our selected time-series offered poor reliability for estimating population trends, consistent with previous studies (Cortés-Guzmán et al., 2026; Duchenne et al., 2022; Leung & Gonzalez, 2024) because they were short and with substantial inter-annual variability. Most of the time-series were expected to provide high estimation error, high false-positive rates, as well as very little power to detect gradual population declines. Many of the analysed BioTime-series had insufficient expected power to detect gradual population declines, preventing us to identify slow long-term declines. However, at least temporal trends are rarely estimated in the wrong direction, as shown for other datasets (Cortés-Guzmán et al., 2026; Wauchope et al., 2019).

Because time-series characteristics are closely linked to the quality of the monitoring but also to species characteristics –life history, detectability in the field, etc.–, different taxa have different levels of expected reliability. Insects showed especially low expected reliability, although a good proportion of their time-series were relatively long. The low expected reliability was partially due to their high inter-annual variability in abundance, likely linked to their life history and population size (Dallas & Kramer, 2022; Montgomery et al., 2026). Such variability makes it difficult to distinguish temporary fluctuations from persistent long-term change. Contrary to previous suggestions that a decade is sufficient for detecting immediate threats in short-lived organisms when data quality is high (Rueda-Cediel et al., 2015), our analyses indicated that reliability is not enough after 10 years (Fig. S7). Interestingly, restricting analyses to insect time-series offering high reliability in estimating population trends (estimation error < 2%.year⁻¹) produced a more negative average trend (−1.35%.year⁻¹, n=689) than the complete dataset (−0.89%.year⁻¹, n = 4385, Table S1). This pattern could be explained because reliable time-series tend to be longer and therefore have an older baseline, accumulating more anthropogenic disturbances (Mihoub et al., 2017; Rodrigues et al., 2019), but might also reflect the fact that most of the estimates of population trends are unreliable and misestimate true insect decline. Consequently, some disagreement among published estimates of insect population changes (Cardinale et al., 2018; Crossley et al., 2020; Simmons et al., 2019), may arise not only from biological or monitoring differences among studies (Duchenne et al., 2022), but also from differences in the reliability offered by the underlying time-series.

Mammals, amphibians, squamates and arachnids showed very low average reliability, mainly because available time-series tended to be short. This limitation was particularly pronounced for amphibians, squamates, and arachnids, highlighting the need for greater investment in long-term monitoring of these comparatively underrepresented taxa (Awkerman et al., 2020; Green, 2003). This was the case even for mammals, for which despite many time-series being available, most were too short to estimate reliable population trends. This is interesting as mammal’s time-series also tended to exhibit negative autocorrelation, a pattern that favours trend estimation. Birds were the only taxonomic group showing substantially higher expected reliability, mainly because it was the group with the longer time-series duration, despite their slightly positive temporal autocorrelation. This reflects how long-term investment in monitoring programs, such as the North American Breeding Bird Survey (Sauer et al., 2017), can improve population trend reliability. Nevertheless, nearly 80% of these bird populations lacked sufficient statistical power to detect gradual declines of 2%.year^-1^, indicating that slow population declines are likely to be undetected.

### Improving reliability in population trends: challenges and perspectives

The main factor that ecologists can manipulate to improve reliability of the estimated population trends is the duration of the monitoring. However, improving reliability of estimated population trends is challenged not only by limited temporal coverage, but also by the inherent stochasticity of ecological systems. While a part of the inter-annual variability and its autocorrelation are intrinsic properties of the population dynamics, there are other sources of noise that can be reduced by researchers by increasing data quality. Reliability is not only “the longer the better”: extending monitoring duration generally improves trend reliability, but improving data quality may sometimes be more effective than simply collecting additional years (Rueda-Cediel et al., 2015). In real monitoring schemes, factors such as opportunistic data collection, unexpected weather issues, problems with funding and time (Cortés-Guzmán et al., 2026), observer identity turnover (Vallecillo et al., 2021), changes in sampling protocols (Chang et al., 2023), and site-selection biases (Fournier et al., 2019) reduce reliability by directly affecting inter-annual variability and its autocorrelation. Consistent sampling protocols, observer training, and taxon-specific monitoring designs can reduce observation-related variability and improve the comparability of long-term biodiversity data (Potts et al., 2024). The use of spatial and phenological high coverage methods, such as camera traps, bioacoustics (Crochard et al., 2026), artificial monitoring landscapes (Ash et al., 2026) or automated monitoring (Besson et al., 2022) could reduce sources of variability due to changing observation biases. Lastly, although we focused on linear population trends using GLMMs, population trajectories are often nonlinear and may require more flexible approaches, such as GAMs. However, the additional parameters required by these models may increase uncertainty and the risk of overfitting, particularly in short and highly variable time-series (Edwards et al., 2025; Schmucki et al., 2016).

Additionally, reliability can be achieved by replicating measures across space (Nielsen et al., 2009). Spatial replication may be more efficient than extending monitoring duration, depending on temporal and spatial autocorrelation (Rhodes & Jonzén, 2011). The less positive the autocorrelation, the more efficient the sampling. Thus, monitoring several sites might be more efficient than extending survey through time, providing a practical route to improve reliability when long time-series are unavailable. Specifically, increasing from one to a dozen sites in our simulated data can significantly increase reliability (Supplementary methods; Fig. S8). When data are multi-taxa, a similar strategy is the use of composite indicators (Collen et al., 2009; Edwards et al., 2025; PECBMS, 2025), although they have been shown to obscure site-specific or species-specific responses and can themselves be sensitive to methodological choices (Toszogyova et al., 2024). Additionally, a complementary strategy could be to recover temporal depth through digitalisation of historical records collected under consistent protocols (Neff et al., 2026) to extend time-series and improve our ability to assess ecological changes.

One of the major challenges in biodiversity conservation is the mismatch between the time scale needed for policy decisions and the time scale required to detect true population changes. Our results show that evaluating conservation interventions short enough can produce misleading conclusions because population trends are unreliable over the short term, leading to delayed actions or misallocating resources (Didham et al., 2020). Most of the recent conservation actions developed at international levels operate on policy cycles of only a few years, such as the EU pollinator monitoring scheme (Potts et al., 2024), the Kunming–Montreal Global Biodiversity Framework (Convention on Biological Diversity, 2022), the Nature Restoration Law (EU, 2024), and the Common Agricultural Policy (EU, 2021). Long-term funding commitments remain essential, as duration consistently emerged as the strongest determinant of reliable inference. Ultimately, achieving reliable biodiversity indicators will require recognizing that robust ecological inference is not only a scientific challenge, but also a long-term institutional commitment.

## Data availability

Empirical time-series are available from BioTIME v2.0: https:/io.st-andrews.ac.uk/. R code to reproduce all analyses and intermediate outputs isavailable at Zenodo https://zenodo.org/records/23268178 and GitHub https://github.com/juliagdealedo/uncertainty_time_series.

## Statement of authorship

JGA, FD, and IB conceived the study. JGA retrieved and cleaned the data. JGA and FD performed the simulations and analysed the data. JGA wrote the first draft of the manuscript with ongoing input from IB and FD, and all authors contributed substantially to its development and revision.

## Acknowledgements

This research was funded by Biodiversa+, the European Biodiversity Partnership, through the ANTENNA project under the 2022–2023 BiodivMon joint call. ANTENNA was co-funded by the European Commission (GA No. 101052342) and the following national and regional funding organisations: Deutsche Forschungsgemeinschaft e.V.; BMBF-VDI/VDE Innovation + Technik GmbH; Dutch Research Council; Innovation Fund Denmark; Agencia Estatal de Investigacion; Fundacion Biodiversidad; General Secretariat for Research and Innovation; and Environmental Protection Agency of Ireland. The Spanish contribution to ANTENNA was supported through PCI2023-146022-2, funded by the Spanish State Research Agency (Agencia Estatal de Investigacion, AEI) and co-funded by the European Union. F.D. was funded by a postdoc Mobility grant from the SNF (project number: P500PB_217801).

## References

Ash, A., Hallett, S., Carvell, C., Williams, L., & August, T. (2026). Testing the efficacy of artificial flowers as a novel attractant for automated pollinator monitoring. Journal of Pollination Ecology, 41, 137–153. 10.26786/1920-7603(2026)893

Awkerman, J., Raimondo, S., Schmolke, A., Galic, N., Rueda-Cediel, P., Kapo, K., Accolla, C., Vaugeois, M., & Forbes, V. (2020). Guidance for Developing Amphibian Population Models for Ecological Risk Assessment. Integrated Environmental Assessment and Management, 16(2), 223–233. 10.1002/ieam.4215

Besson, M., Alison, J., Bjerge, K., Gorochowski, T. E., Høye, T. T., Jucker, T., Mann, H. M. R., & Clements, C. F. (2022). Towards the fully automated monitoring of ecological communities. Ecology Letters, 25(12), 2753–2775. 10.1111/ele.14123

Cardinale, B. J., Gonzalez, A., Allington, G. R. H., & Loreau, M. (2018). Is local biodiversity declining or not? A summary of the debate over analysis of species richness time trends. Biological Conservation, 219, 175–183. 10.1016/j.biocon.2017.12.021

Ceballos, G., & Ehrlich, P. R. (2023). Mutilation of the tree of life via mass extinction of animal genera. Proceedings of the National Academy of Sciences, 120(39), e2306987120. 10.1073/pnas.2306987120

Ceballos, G., Ehrlich, P. R., & Dirzo, R. (2017). Biological annihilation via the ongoing sixth mass extinction signaled by vertebrate population losses and declines. Proceedings of the National Academy of Sciences, 114(30), E6089–E6096. 10.1073/pnas.1704949114

Chaikin, S., Riva, F., Marshall, K. E., Lessard, J.-P., & Belmaker, J. (2024). Marine fishes experiencing high-velocity range shifts may not be climate change winners. Nature Ecology & Evolution, 8(5), 936–946. 10.1038/s41559-024-02350-7

Chamberlain, S., Barve, V., Mcglinn, D., Oldoni, D., Desmet, P., Geffert, L., & Ram, K. (2025). rgbif: Interface to the Global Biodiversity Information Facility API. https://CRAN.R-project.org/package=rgbif

Chang, H.-Y., Sun, M., Rokosz, K., & Chen, Y. (2023). Evaluating effects of changing sampling protocol for a long-term ichthyoplankton monitoring program. Frontiers in Marine Science, 10. 10.3389/fmars.2023.1237549

Cohen, J. (1988). Statistical power analysis for the behavioral sciences (2nd ed). L. Erlbaum Associates.

Collen, B., Loh, J., Whitmee, S., McRAE, L., Amin, R., & Baillie, J. E. M. (2009). Monitoring Change in Vertebrate Abundance: The Living Planet Index. Conservation Biology, 23(2), 317–327. 10.1111/j.1523-1739.2008.01117.x

Connors, B. M., Cooper, A. B., Peterman, R. M., & Dulvy, N. K. (2014). The false classification of extinction risk in noisy environments. Proceedings of the Royal Society B: Biological Sciences, 281(1787), 20132935. 10.1098/rspb.2013.2935

Convention on Biological Diversity. (2022). Kunming-Montreal Global Biodiversity Framework (15th meeting-Part II). Convention on Biological Diversity. https://www.cbd.int/doc/decisions/cop-15/cop-15-dec-04-en.pdf

Cortés-Guzmán, D., Sinclair, J. S., Aroviita, J., Azpiroz, I., de Baat, M. L., Bañares, I., Becker, E., Cañedo-Argüelles, M., Cosson, E., Cunillera-Montcusí, D., Escaffre, R., Ferréol, M., Forio, M. A. E., Goethals, P., González-Ferreras, A. M., Huttunen, K.-L., Larrañaga, A., López, E. S., Rubio, M., … Haase, P. (2026). Robust estimates of biodiversity change require high-resolution time-series. Nature Communications, 17(1), 5941. 10.1038/s41467-026-75321-0

Crochard, L., Mariton, L., Fontaine, C., Baude, M., Ragué, M., Bas, D., Gaba, S., Bretagnolle, V., Julliard, R., & Bas, Y. (2026). Buzzy bees: Improving the monitoring of pollinator activity in sunflower fields with continuous acoustic recording and deep learning. Ecological Informatics, 94, 103633. 10.1016/j.ecoinf.2026.103633

Crossley, M. S., Meier, A. R., Baldwin, E. M., Berry, L. L., Crenshaw, L. C., Hartman, G. L., Lagos-Kutz, D., Nichols, D. H., Patel, K., Varriano, S., Snyder, W. E., & Moran, M. D. (2020). No net insect abundance and diversity declines across US Long Term Ecological Research sites. Nature Ecology & Evolution, 4(10), 1368–1376. 10.1038/s41559-020-1269-4

Dallas, T. A., & Kramer, A. M. (2022). Temporal variability in population and community dynamics. Ecology, 103(2), e03577. 10.1002/ecy.3577

Daskalova, G. N., Myers-Smith, I. H., Bjorkman, A. D., Blowes, S. A., Supp, S. R., Magurran, A. E., & Dornelas, M. (2020). Landscape-scale forest loss as a catalyst of population and biodiversity change. Science, 368(6497), 1341– 1347. 10.1126/science.aba1289

Daskalova, G. N., Phillimore, A. B., & Myers-Smith, I. H. (2021). Accounting for year effects and sampling error in temporal analyses of invertebrate population and biodiversity change: A comment on Seibold et al. 2019. Insect Conservation and Diversity, 14(1), 149–154. 10.1111/icad.12468

Dennis, E. B., Morgan, B. J. T., Freeman, S. N., Brereton, T. M., & Roy, D. B. (2016). A generalized abundance index for seasonal invertebrates. Biometrics, 72(4), 1305–1314. 10.1111/biom.12506

Didham, R. K., Basset, Y., Collins, C. M., Leather, S. R., Littlewood, N. A., Menz, M. H. M., Müller, J., Packer, L., Saunders, M. E., Schönrogge, K., Stewart, A. J. A., Yanoviak, S. P., & Hassall, C. (2020). Interpreting insect declines: Seven challenges and a way forward. Insect Conservation and Diversity, 13(2), 103–114. 10.1111/icad.12408

Dirzo, R., & Raven, P. H. (2003). Global state of biodiversity and loss. Annual Review of Environment and Resources, 28(1), 137–167.

Dornelas, M., Antão, L. H., Bates, A. E., Brambilla, V., Chase, J. M., Chow, C. F. Y., Fontrodona-Eslava, A., Magurran, A. E., Martins, I. S., Moyes, F., Sagouis, A., Adu-Acheampong, S., Acquah-Lamptey, D., Adam, D., Ajani, P. A., Albaina, A., Almaraz, P., An, J., Anderson, R. S., … Zvuloni, A. (2025). BioTIME 2.0: Expanding and Improving a Database of Biodiversity Time-series. Global Ecology and Biogeography, 34(5), e70003. 10.1111/geb.70003

Dornelas, M., Antão, L. H., Moyes, F., Bates, A. E., Magurran, A. E., Adam, D., Akhmetzhanova, A. A., Appeltans, W., Arcos, J. M., Arnold, H., Ayyappan, N., Badihi, G., Baird, A. H., Barbosa, M., Barreto, T. E., Bässler, C., Bellgrove, A., Belmaker, J., Benedetti-Cecchi, L., … Zettler, M. L. (2018). BioTIME: A database of biodiversity time-series for the Anthropocene. Global Ecology and Biogeography, 27(7), 760–786. 10.1111/geb.12729

Dornelas, M., Gotelli, N. J., McGill, B., Shimadzu, H., Moyes, F., Sievers, C., & Magurran, A. E. (2014). Assemblage time-series reveal biodiversity change but not systematic loss. 10.1126/science.1248484

Dove, S., Böhm, M., Freeman, R., McRae, L., & Murrell, D. J. (2023). Quantifying reliability and data deficiency in global vertebrate population trends using the Living Planet Index. Global Change Biology, 29(17), 4966–4982. 10.1111/gcb.16841

Duchenne, F., Porcher, E., Mihoub, J.-B., Loïs, G., & Fontaine, C. (2022). Controversy over the decline of arthropods: A matter of temporal baseline? Peer Community Journal, 2. 10.24072/pcjournal.131

Edwards, C. B., & Crone, E. E. (2021). Estimating abundance and phenology from transect count data with GLMs. Oikos, 130(8), 1335–1345. 10.1111/oik.08368

Edwards, C. B., Zipkin, E. F., Henry, E. H., Haddad, N. M., Forister, M. L., Burls, K. J., Campbell, S. P., Crone, E. E., Diffendorfer, J., Douglas, M. R., Drum, R. G., Fallon, C. E., Glassberg, J., Grames, E. M., Hatfield, R., Hershcovich, S., Hoffman Black, S., Larsen, E. A., Leuenberger, W., … Schultz, C. B. (2025). Rapid butterfly declines across the United States during the 21st century. Science, 387(6738), 1090–1094. 10.1126/science.adp4671

Engen, S., Sæther, B.-E., Armitage, K. B., Blumstein, D. T., Clutton-Brock, T. H., Dobson, F. S., Festa-Bianchet, M., Oli, M. K., & Ozgul, A. (2013). Estimating the effect of temporally autocorrelated environments on the demography of density-independent age-structured populations. Methods in Ecology and Evolution, 4(6), 573–584. 10.1111/2041-210X.12043

Fournier, A. M. V., White, E. R., & Heard, S. B. (2019). Site-selection bias and apparent population declines in long-term studies. Conservation Biology, 33(6), 1370–1379. 10.1111/cobi.13371

Gaume, L., & Desquilbet, M. (2024). InsectChange: Comment. Peer Community Journal, 4. 10.24072/pcjournal.469

Gelman, A., & Carlin, J. (2014). Beyond Power Calculations: Assessing Type S (Sign) and Type M (Magnitude) Errors. Perspectives on Psychological Science, 9(6), 641–651. 10.1177/1745691614551642

Gerrodette, T. (1987). A Power Analysis for Detecting Trends. Ecology, 68(5), 1364– 1372. 10.2307/1939220

Green, D. M. (2003). The ecology of extinction: Population fluctuation and decline in amphibians. Biological Conservation, 111(3), 331–343. 10.1016/S0006-3207(02)00302-6

Hansen, T. F., Stenseth, N. C., Henttonen, H., & Tast, J. (1999). Interspecific and intraspecific competition as causes of direct and delayed density dependence in a fluctuating vole population. Proceedings of the National Academy of Sciences, 96(3), 986–991. 10.1073/pnas.96.3.986

Jaureguiberry, P., Titeux, N., Wiemers, M., Bowler, D. E., Coscieme, L., Golden, A. S., Guerra, C. A., Jacob, U., Takahashi, Y., Settele, J., Díaz, S., Molnár, Z., & Purvis, A. (2022). The direct drivers of recent global anthropogenic biodiversity loss. Science Advances, 8(45), eabm9982. 10.1126/sciadv.abm9982

Johnson, P. C. D., Barry, S. J. E., Ferguson, H. M., & Müller, P. (2015). Power analysis for generalized linear mixed models in ecology and evolution. Methods in Ecology and Evolution, 6(2), 133–142. 10.1111/2041-210X.12306

Johnson, T. F., Beckerman, A. P., Childs, D. Z., Webb, T. J., Evans, K. L., Griffiths, C. A., Capdevila, P., Clements, C. F., Besson, M., Gregory, R. D., Thomas, G. H., Delmas, E., & Freckleton, R. P. (2024). Revealing uncertainty in the status of biodiversity change. Nature, 628(8009), 788–794. 10.1038/s41586-024-07236-z

Justin Nowakowski, A., Watling, J. I., Murray, A., Deichmann, J. L., Akre, T. S., Muñoz Brenes, C. L., Todd, B. D., McRae, L., Freeman, R., & Frishkoff, L. O. (2023). Protected areas slow declines unevenly across the tetrapod tree of life. Nature, 622(7981), 101–106. 10.1038/s41586-023-06562-y

Leung, B., & Gonzalez, A. (2024). Global monitoring for biodiversity: Uncertainty, risk, and power analyses to support trend change detection. Science Advances, 10(7), eadj1448. 10.1126/sciadv.adj1448

Loreau, M., Cardinale, B. J., Isbell, F., Newbold, T., O’Connor, M. I., & de Mazancourt, C. (2022). Do not downplay biodiversity loss. Nature, 601(7894), E27–E28. 10.1038/s41586-021-04179-7

Mihoub, J.-B., Henle, K., Titeux, N., Brotons, L., Brummitt, N. A., & Schmeller, D. S. (2017). Setting temporal baselines for biodiversity: The limits of available monitoring data for capturing the full impact of anthropogenic pressures. Scientific Reports, 7. 10.1038/srep41591

Montgomery, G. A., Grames, E. M., Jacobsen, J. M., Kahn, E. X., Leyel, A., Rosati, M. C., Wu, Q., Elphick, C. S., Wagner, D., & Tingley, M. W. (2026). Are Insect Populations Inherently More Variable? A Multi-Taxa Approach to Characterising Interannual Fluctuations in Insect Time-series. Ecology Letters, 29(3), e70359. 10.1111/ele.70359

Murphy, G. E. P., & Romanuk, T. N. (2014). A meta-analysis of declines in local species richness from human disturbances. Ecology and Evolution, 4(1), 91– 103. 10.1002/ece3.909

Neff, F., Bollmann, K., Chittaro, Y., Gossner, M. M., Herzog, F., Korner-Nievergelt, F., Litsios, G., Martínez-Núñez, C., Moretti, M., Rey, E., Sanchez, A., & Knop, E. (2026). Ninety-year trends reveal sharpest insect declines in the mid-twentieth century. Nature Ecology & Evolution, 1–11. 10.1038/s41559-026-03074-6

Nielsen, S. E., Haughland, D. L., Bayne, E., & Schieck, J. (2009). Capacity of large-scale, long-term biodiversity monitoring programmes to detect trends in species prevalence. Biodiversity and Conservation, 18(11), 2961–2978. 10.1007/s10531-009-9619-1

Nicolau, P. G., Ims, R. A., Sørbye, S. H., & Yoccoz, N. G. (2022). Seasonality, density dependence, and spatial population synchrony. Proceedings of the National Academy of Sciences, 119(51), e2210144119. 10.1073/pnas.2210144119

PECBMS. (2025, December 15). European Wild Bird Indices, 2025 update. PECBMS. https://pecbms.info/european-wild-bird-indices-2025-update/

Pimm, S., Jenkins, C., Abell, R., Brooks, T., Gittleman, J., Joppa, L., Raven, P., Roberts, C., & Sexton, J. (2014). The biodiversity of species and their rates of extinction, distribution, and protection. Science, 344, 1246752. 10.1126/science.1246752

Potts, S. G., Bartomeus, I., Biesmeijer, K., Breeze, T., Casino, A., Dauber, J., Dieker, P., Hochkirch, A., Høye, T., Isaac, N., Kleijn, D., Laikre, L., Mandelik, Y., Montagna, M., Montero Castaño, A., Öckinger, E., Oteman, B., Pardo Valle, A., Polce, C., … Zhang, J. (2024). Refined proposal for an EU pollinator monitoring scheme. European Union. Publications Office. 10.2760/2005545

Regulation (EU) 2024/1991 of the European Parliament and of the Council of 24 June 2024 on Nature Restoration and Amending Regulation (EU) 2022/869 (Text with EEA Relevance) (2024). http://data.europa.eu/eli/reg/2024/1991/oj

Regulation (EU) 2021/2115 of the European Parliament and of the Council of 2 December 2021 Establishing Rules on Support for Strategic Plans to Be Drawn up by Member States under the Common Agricultural Policy (CAP Strategic Plans) and Financed by the European Agricultural Guarantee Fund (EAGF) and by the European Agricultural Fund for Rural Development (EAFRD) and Repealing Regulations (EU) No 1305/2013 and (EU) No 1307/2013, 435 OJ L (2021). http://data.europa.eu/eli/reg/2021/2115/oj

Rhodes, J. R., & Jonzén, N. (2011). Monitoring temporal trends in spatially structured populations: How should sampling effort be allocated between space and time? Ecography, 34(6), 1040–1048. 10.1111/j.1600-0587.2011.06370.x

Rodrigues, A. S. L., Monsarrat, S., Charpentier, A., Brooks, T. M., Hoffmann, M., Reeves, R., Palomares, M. L. D., & Turvey, S. T. (2019). Unshifting the baseline: A framework for documenting historical population changes and assessing long-term anthropogenic impacts. Philosophical Transactions of the Royal Society B: Biological Sciences, 374(1788), 20190220. 10.1098/rstb.2019.0220

Rueda-Cediel, P., Anderson, K. E., Regan, T. J., Franklin, J., & Regan, H. M. (2015). Combined Influences of Model Choice, Data Quality, and Data Quantity When Estimating Population Trends. PLOS ONE, 10(7), e0132255. 10.1371/journal.pone.0132255

Sala, O. E., Chapin, F. S., Iii Armesto, J. J., Berlow, E., Bloomfield, J., Dirzo, R., Huber-Sanwald, E., Huenneke, L. F., Jackson, R. B., Kinzig, A., Leemans, R., Lodge, D. M., Mooney, H. A., Oesterheld, M., Poff, N. L., Sykes, M. T., Walker, B. H., Walker, M., & Wall, D. H. (2000). Global Biodiversity Scenarios for the Year 2100. Science, 287(5459), 1770–1774. 10.1126/science.287.5459.1770

Sauer, J. R., Pardieck, K. L., Ziolkowski, D. J., Smith, A. C., Hudson, M.-A. R., Rodriguez, V., Berlanga, H., Niven, D. K., & Link, W. A. (2017). The first 50 years of the North American Breeding Bird Survey. The Condor, 119(3), 576– 593. 10.1650/CONDOR-17-83.1

Schmucki, R., Pe’er, G., Roy, D. B., Stefanescu, C., Van Swaay, C. A. M., Oliver, T. H., Kuussaari, M., Van Strien, A. J., Ries, L., Settele, J., Musche, M., Carnicer, J., Schweiger, O., Brereton, T. M., Harpke, A., Heliölä, J., Kühn, E., & Julliard, R. (2016). A regionally informed abundance index for supporting integrative analyses across butterfly monitoring schemes. Journal of Applied Ecology, 53(2), 501–510. 10.1111/1365-2664.12561

Simmons, B. I., Balmford, A., Bladon, A. J., Christie, A. P., De Palma, A., Dicks, L. V., Gallego-Zamorano, J., Johnston, A., Martin, P. A., Purvis, A., Rocha, R., Wauchope, H. S., Wordley, C. F. R., Worthington, T. A., & Finch, T. (2019). Worldwide insect declines: An important message, but interpret with caution. Ecology and Evolution, 9(7), 3678–3680. 10.1002/ece3.5153

Steidl, R. J., & Thomas, L. (2001). Power Analysis and Experimental Design. In S. M. Scheiner & J. Gurevitch (Eds.), Design and Analysis of Ecological Experiments (p. 0). Oxford University Press. 10.1093/oso/9780195131871.003.0002

Toszogyova, A., Smyčka, J., & Storch, D. (2024). Mathematical biases in the calculation of the Living Planet Index lead to overestimation of vertebrate population decline. Nature Communications, 15(1), 5295. 10.1038/s41467-024-49070-x

Valdez, J. W., Callaghan, C. T., Junker, J., Purvis, A., Hill, S. L. L., & Pereira, H. M. (2023). The undetectability of global biodiversity trends using local species richness. Ecography, 2023(3), e06604. 10.1111/ecog.06604

Vallecillo, D., Gauthier-Clerc, M., Guillemain, M., Vittecoq, M., Vandewalle, P., Roche, B., & Champagnon, J. (2021). Reliability of animal counts and implications for the interpretation of trends. Ecology and Evolution, 11(5), 2249–2260. 10.1002/ece3.7191

Vindenes, Y., & Engen, S. (2017). Demographic stochasticity and temporal autocorrelation in the dynamics of structured populations. Oikos, 126(4), 462– 471. 10.1111/oik.03958

Wauchope, H. S., Amano, T., Sutherland, W. J., & Johnston, A. (2019). When can we trust population trends? A method for quantifying the effects of sampling interval and duration. Methods in Ecology and Evolution, 10(12), 2067–2078. 10.1111/2041-210X.13302

White, E. R. (2019). Minimum Time Required to Detect Population Trends: The Need for Long-Term Monitoring Programs. BioScience, 69(1), 40–46. 10.1093/biosci/biy144

Wilson, H. B., Kendall, B. E., & Possingham, H. P. (2011). Variability in Population Abundance and the Classification of Extinction Risk. Conservation Biology, 25(4), 747–757. 10.1111/j.1523-1739.2011.01671.x

Yang, Y., Hillebrand, H., Lagisz, M., Cleasby, I., & Nakagawa, S. (2022). Low statistical power and overestimated anthropogenic impacts, exacerbated by publication bias, dominate field studies in global change biology. Global Change Biology, 28(3), 969–989. 10.1111/gcb.15972

