## Supplementary material for "Available time-series of terrestrial animals provide low reliability in population trend estimates"

**Available time series provide low reliability in estimating population trends for terrestrial animals**

Content

### Supplementary methods

##### Testing the effect of partial-pooling in reducing error rates

Analysing multiple time series simultaneously can increase the power of the analysis. For example, this is done when the same species is monitored on multiple sites or to calculate aggregated trends across species (e.g. Living Planet Index (Collen et al., 2009), Wild Bird Indicators (Gregory & Strien, 2010)). Despite a full analysis on aggregation methods is beyond the scope of this paper, it is illustrative to show power gains as we increase the number of time series simultaneously analysed. To that end, as a proof of concept, we select among our simulated time series those with 10 years and no autocorrelation for two inter-annual variability levels (0.15 and 1.5), and test how the error rate is reduced as we increase the number of time series jointly modelled. To model multiple time series, we use partial pooling by adding to the model structure simulation Id as random intercept. We randomly pool from 1 to 50 simulated time-series and calculated the error between the estimated and true trend. The real trend was calculated as the geometric mean of the true trends of all drawn simulations. This process is repeated 100 times per inter-annual variability level (100 replicates × 10 pooling categories × 2 variability levels = 2,000 simulations). Results are presented in Figure S7. Note that here we assumed time series are independent, and hence, the effect size of replication cannot directly be extrapolated to real conditions. In nature pooling by sites will likely show spatial autocorrelation, and pooling by taxon will likely show phylogenetically shared responses. We expect that the positive effect of partial pooling on reliability would be less important if considering spatial/phylogenetic autocorrelation because it would decrease the level of effective replication. However, pooling should anyway help to increase reliability in real situations. To maintain simulations under a reasonable complexity we decided to consider only independent replicates.

### Supplementary Figures

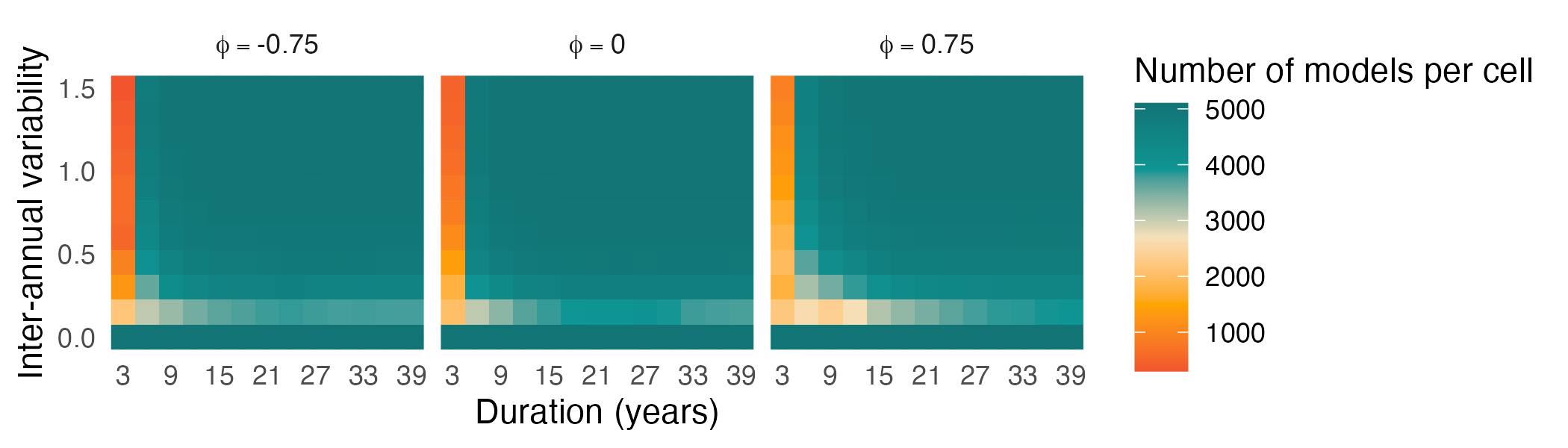

**Fig. S1. Distribution of simulation replicates across the parameter space.** Number of simulated models available for each combination of time-series duration and interannual variability, shown separately for three temporal autocorrelation scenarios (φ = -0.75, 0, and 0.75). Cell values indicate the number of models contributing to each parameter combination and therefore the sample size underlying the reliability metrics estimated in the simulations.

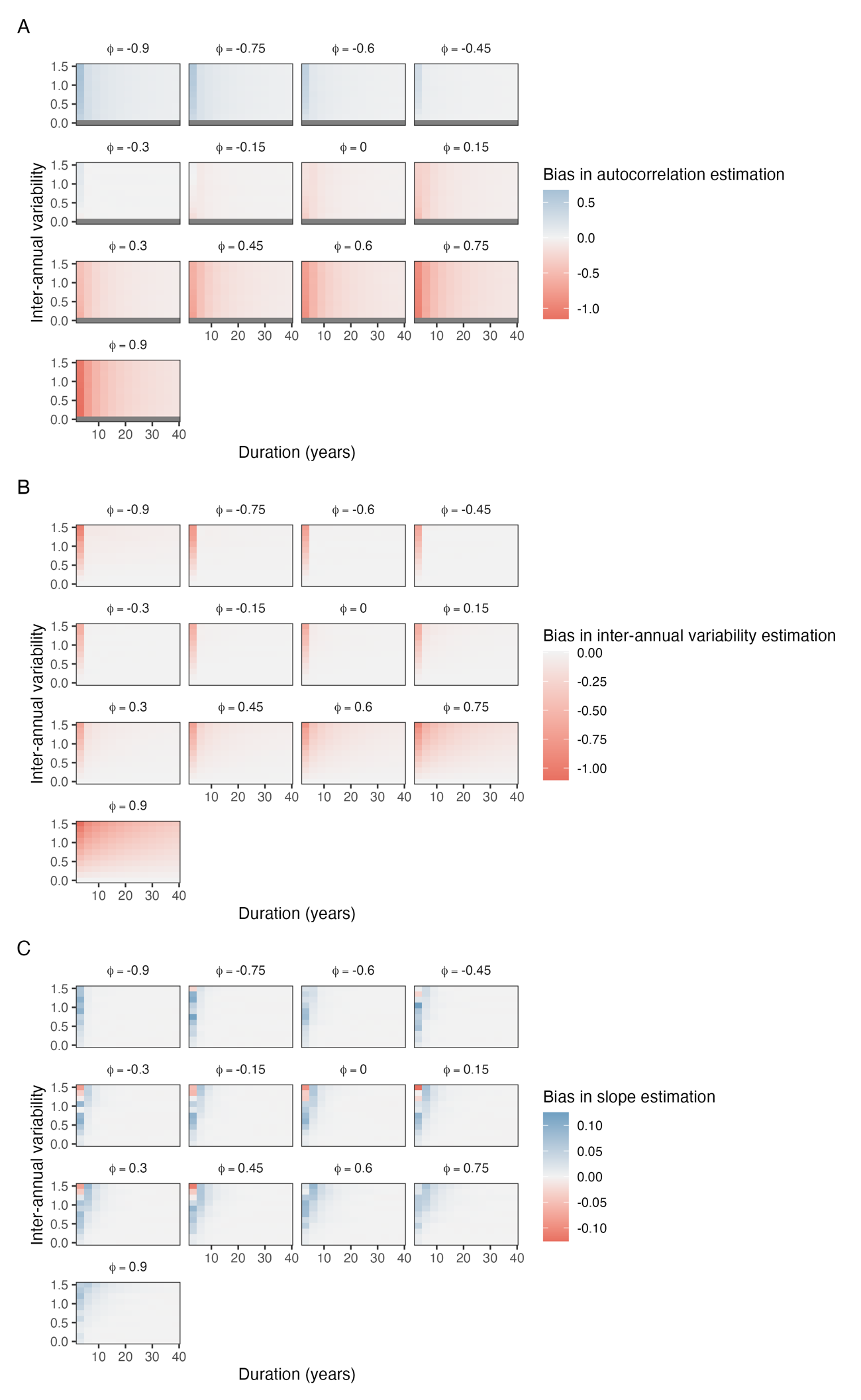

**Fig. S2.** Mean estimation bias for **(A)** temporal autocorrelation (Φ), **(B)** inter-annual variability, and **(C)** temporal trend (slope) across all simulated combinations of monitoring duration, inter-annual variability, and temporal autocorrelation. Heatmaps represent the average difference between estimated and true parameter values, with red colours indicating overestimation and blue colours indicating underestimation. Gray cells represent cells without values, since autocorrelation could not be estimated when not simulating inter-annual variability.

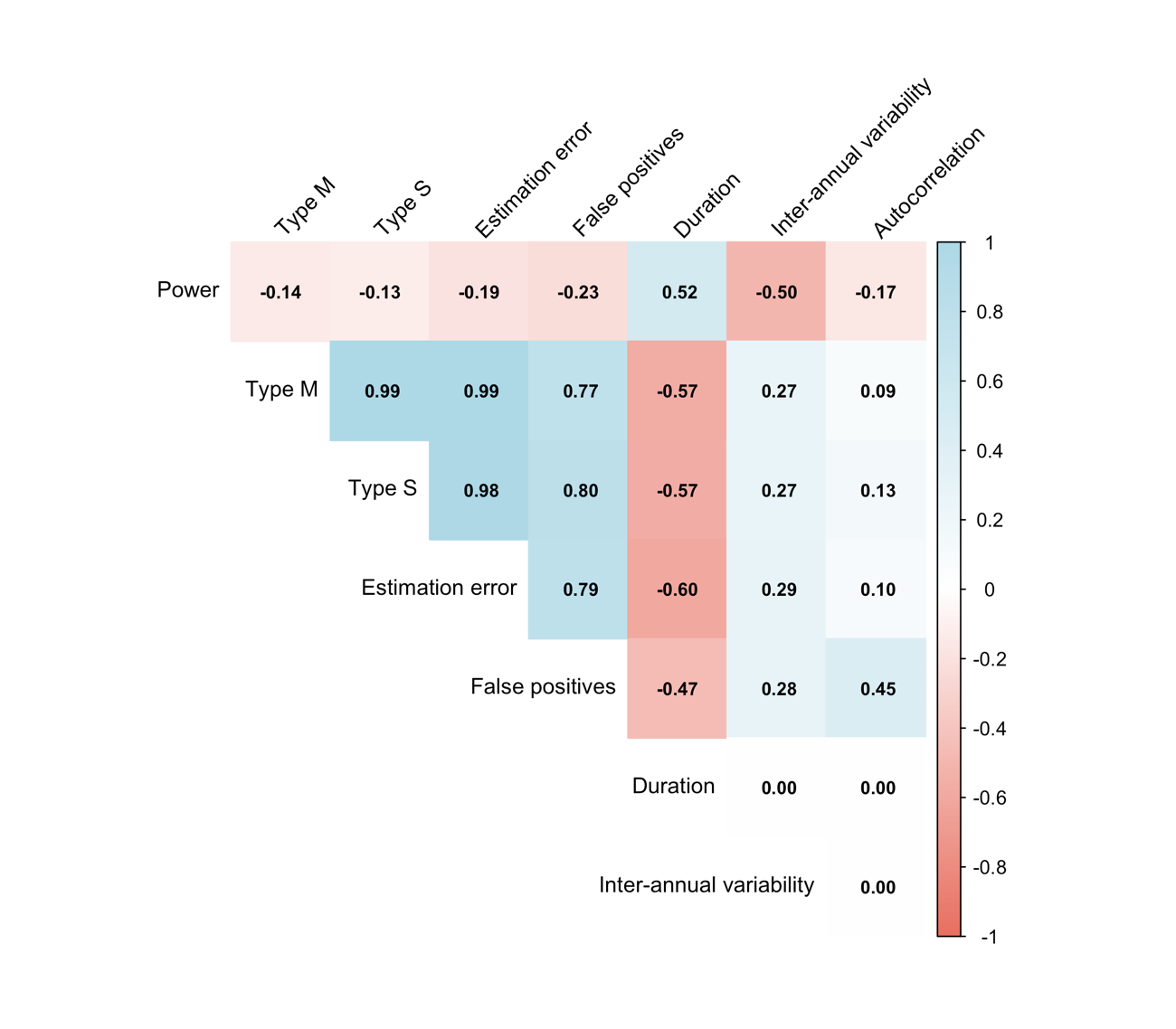

**Fig. S3.** Correlations among trend reliability metrics and time-series characteristics. Pearson correlation coefficients among the five reliability metrics (Type M error, Type S error, estimation error, false-positive rate and power to detect 2%.year^-1^ decline) and the three simulated time-series characteristics (duration, inter-annual variability, and temporal autocorrelation). Values within cells indicate Pearson’s correlation coefficient (r), with colours representing the direction and strength of the association (red, negative; blue, positive). The three time-series characteristics were uncorrelated by design.

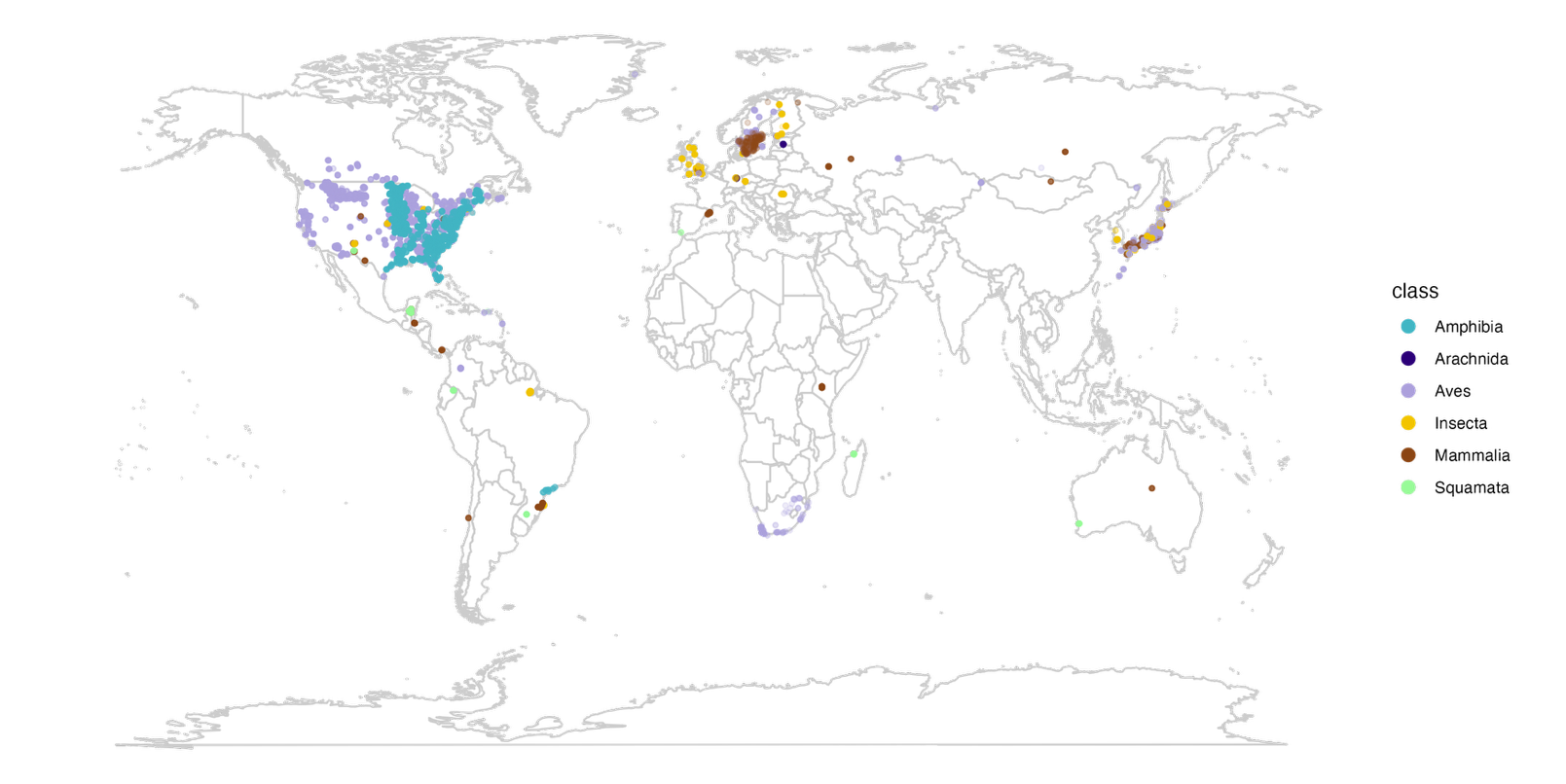

**Fig. S4**. Geographic distribution of empirical population time series used in this study extracted from the BioTIME 2.0 database. Each point represents a population time series and is coloured according to its taxonomic class (Amphibia, Arachnida, Aves, Insecta, Mammalia, and Squamata). Country boundaries are shown for reference.

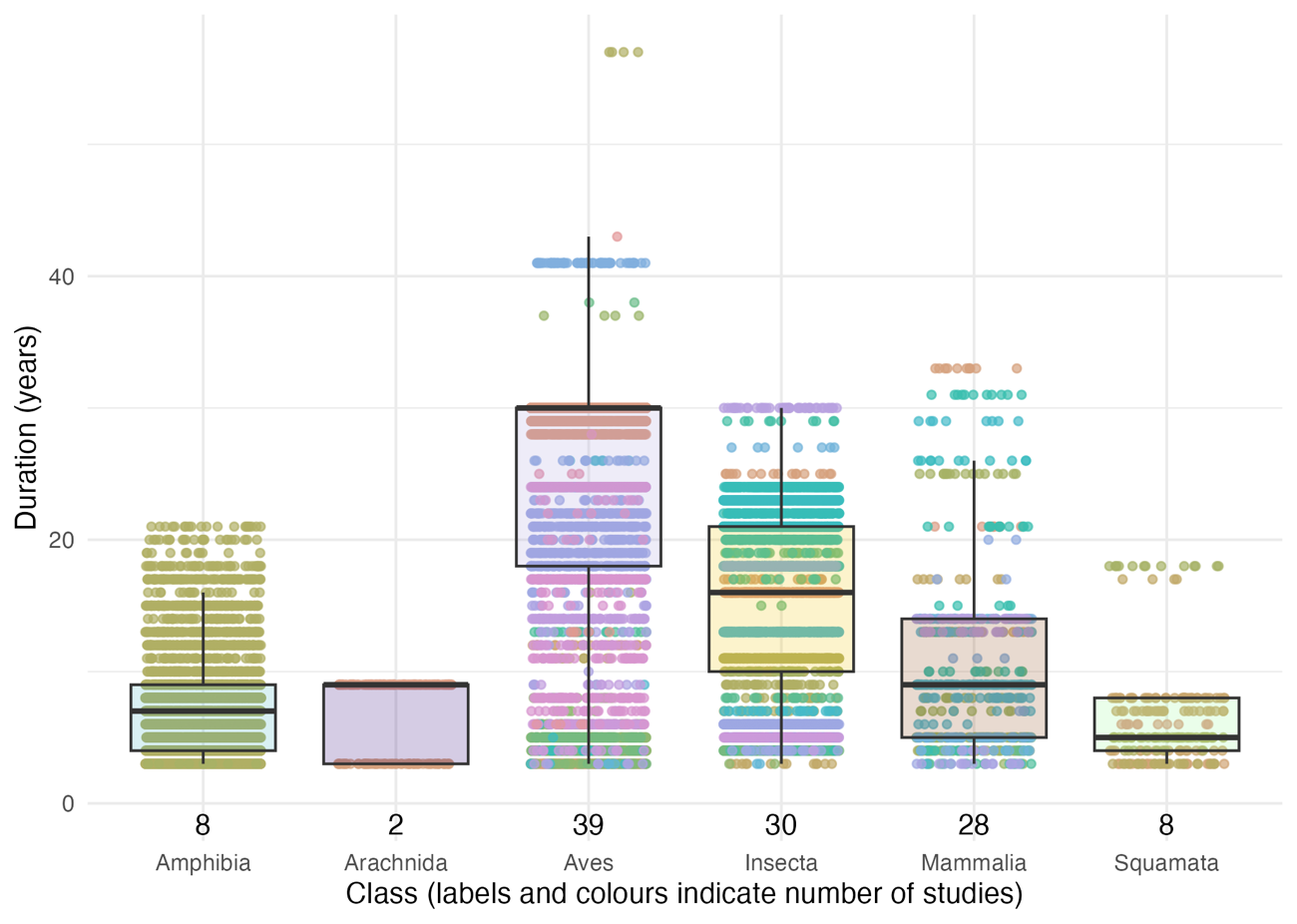

**Fig. S5.** Distribution of time-series duration across taxonomic classes in the BioTIME 2.0 dataset. Each point represents an individual population time series, coloured according to the study from which it originated. Boxplots summarise the median and interquartile range of time-series duration within each class. Numbers above the x-axis indicate the number of independent studies contributing data to each taxonomic group.

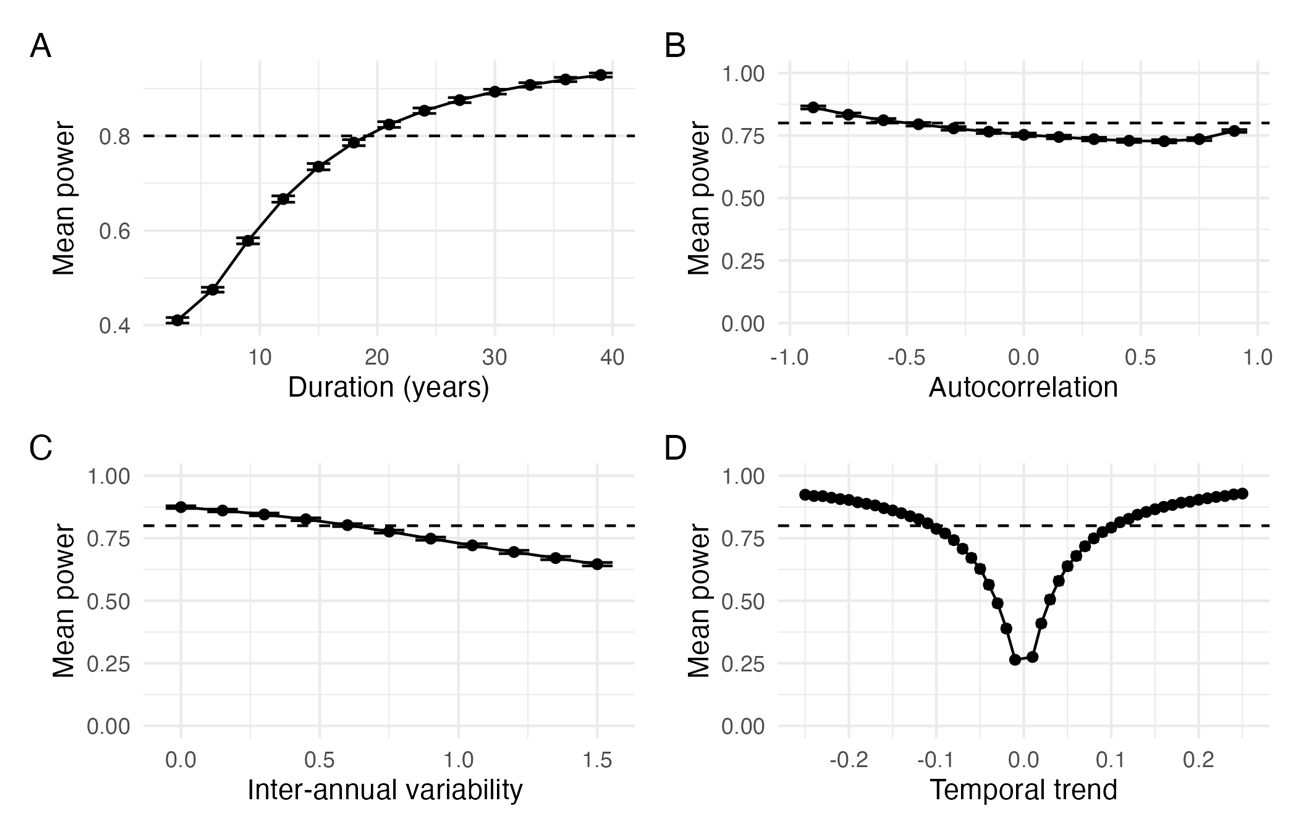

**Fig. S6. Relationship between statistical power and time-series characteristics.** Mean statistical power across simulations as a function of **(A)** duration, **(B)** temporal autocorrelation (Φ), **(C)** interannual variability (CV) and **(D)** annual population trend. Values are averaged across all other simulated parameters. The dashed horizontal line indicates the conventional 80% power threshold. Error bars indicate 95% confidence intervals.

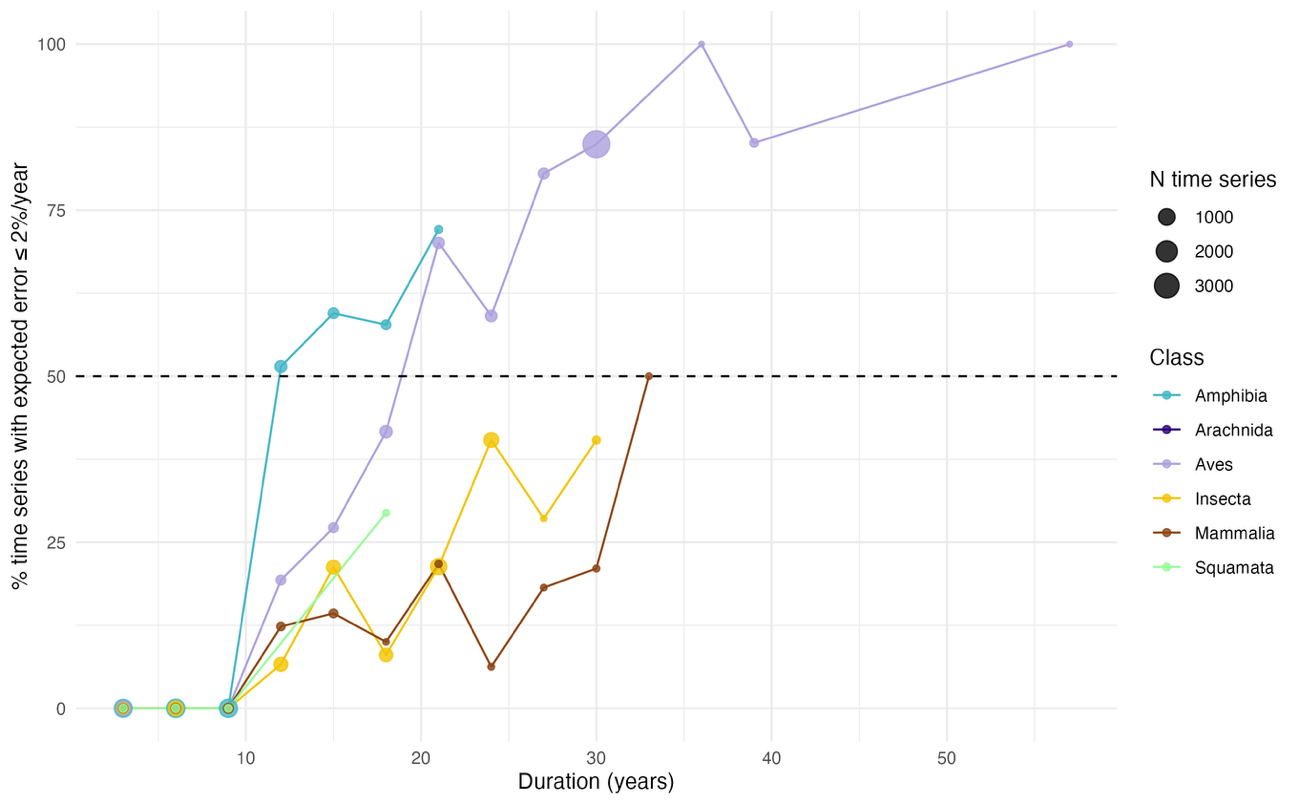

**Fig. S7.** Relationship between time series duration and expected trend reliability in empirical datasets. Percentage of empirical time series with an expected trend estimation error below 2% year⁻¹ as a function of time-series duration. Colours indicate taxonomic class, and point size is proportional to the number of time series available for each duration-class combination. The figure illustrates how the proportion of reliable trend estimates increases with monitoring duration across taxa.

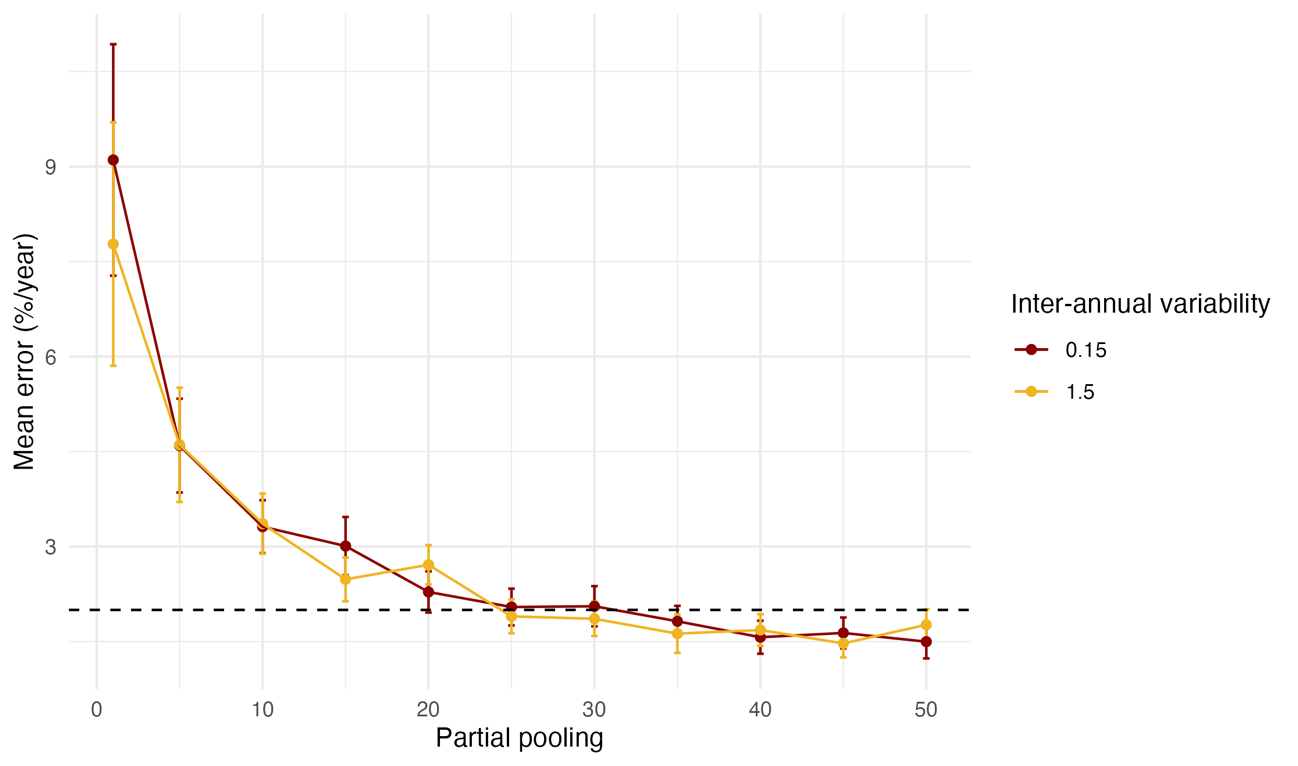

**Fig. S8**. Effect of spatial aggregation on the reliability of trend estimation. Mean error in estimated population trends (% year⁻¹) when abundance time series were aggregated across an increasing number of sites, from 1 to 50. Colours represent the two simulated levels of inter-annual variability (CV = 0.15 and 1.5). Points represent mean estimation error and error bars indicate 95% confidence intervals. The horizontal dashed line indicates the predefined acceptable error threshold of 2% year⁻¹.

| Class | Metric | No. reliable | No. unreliable | No. all | Mean realiable | Mean unreliable | Mean all | Median reliable | Median unreliable | Median all |
| --- | --- | --- | --- | --- | --- | --- | --- | --- | --- | --- |
| **Amphibia** | error | 405 | 4,152 | 4,557 | 0.0031 | 0.0025 | 0.0025 | 0.0048 | 0.0000 | 0.0000 |
|  | false post. | 2,110 | 2,447 | 4,557 | 0.0279 | -0.0194 | 0.0025 | 0.0060 | -0.0057 | 0.0000 |
|  | power | 5 | 4,552 | 4,557 | 0.0377 | 0.0025 | 0.0025 | 0.0161 | 0.0000 | 0.0000 |
|  | typeM | 3,647 | 910 | 4,557 | 0.0025 | 0.0027 | 0.0025 | 0.0000 | 0.0055 | 0.0000 |
|  | typeS | 2,345 | 2,212 | 4,557 | 0.0029 | 0.0021 | 0.0025 | 0.0000 | 0.0000 | 0.0000 |
| **Arachnida** | error |  | 193 | 193 |  | 0.0458 | 0.0458 |  | 0.0218 | 0.0218 |
|  | false post. | 72 | 121 | 193 | 0.0360 | 0.0516 | 0.0458 | 0.0000 | 0.0278 | 0.0218 |
|  | power |  | 193 | 193 |  | 0.0458 | 0.0458 |  | 0.0218 | 0.0218 |
|  | typeM | 186 | 7 | 193 | 0.0463 | 0.0334 | 0.0458 | 0.0212 | 0.0231 | 0.0218 |
|  | typeS | 60 | 133 | 193 | 0.0400 | 0.0484 | 0.0458 | 0.0038 | 0.0232 | 0.0218 |
| **Aves** | error | 4,045 | 2,434 | 6,479 | 0.0007 | 0.0101 | 0.0042 | 0.0010 | 0.0094 | 0.0027 |
|  | false post. | 1,293 | 5,186 | 6,479 | -0.0021 | 0.0058 | 0.0042 | 0.0000 | 0.0042 | 0.0027 |
|  | power | 1,310 | 5,169 | 6,479 | -0.0011 | 0.0056 | 0.0042 | -0.0009 | 0.0046 | 0.0027 |
|  | typeM | 2,159 | 4,320 | 6,479 | 0.0099 | 0.0014 | 0.0042 | 0.0102 | 0.0015 | 0.0027 |
|  | typeS | 5,172 | 1,307 | 6,479 | 0.0022 | 0.0123 | 0.0042 | 0.0020 | 0.0156 | 0.0027 |
| **Insecta** | error | 689 | 3,696 | 4,385 | -0.0135 | -0.0080 | -0.0089 | -0.0159 | -0.0067 | -0.0096 |
|  | false post. | 401 | 3,984 | 4,385 | -0.0228 | -0.0075 | -0.0089 | -0.0193 | -0.0087 | -0.0096 |
|  | power | 28 | 4,357 | 4,385 | -0.0718 | -0.0085 | -0.0089 | -0.0167 | -0.0094 | -0.0096 |
|  | typeM | 945 | 3,440 | 4,385 | -0.0142 | -0.0074 | -0.0089 | -0.0048 | -0.0159 | -0.0096 |
|  | typeS | 2,553 | 1,832 | 4,385 | -0.0157 | 0.0007 | -0.0089 | -0.0139 | 0.0008 | -0.0096 |
| **Mammalia** | error | 39 | 616 | 655 | 0.0280 | 0.0067 | 0.0080 | 0.0081 | 0.0158 | 0.0154 |
|  | false post. | 131 | 524 | 655 | -0.0119 | 0.0130 | 0.0080 | 0.0000 | 0.0160 | 0.0154 |
|  | power | 3 | 652 | 655 | 0.0281 | 0.0079 | 0.0080 | 0.0113 | 0.0155 | 0.0154 |
|  | typeM | 559 | 96 | 655 | 0.0064 | 0.0171 | 0.0080 | 0.0207 | 0.0051 | 0.0154 |
|  | typeS | 277 | 378 | 655 | 0.0171 | 0.0013 | 0.0080 | 0.0087 | 0.0240 | 0.0154 |
| **Squamata** | error | 5 | 191 | 196 | -0.0520 | 0.0239 | 0.0220 | -0.0021 | 0.0889 | 0.0811 |
|  | false post. | 87 | 109 | 196 | -0.0666 | 0.0927 | 0.0220 | 0.0000 | 0.1073 | 0.0811 |
|  | power |  | 196 | 196 |  | 0.0220 | 0.0220 |  | 0.0811 | 0.0811 |
|  | typeM | 179 | 17 | 196 | 0.0058 | 0.1925 | 0.0220 | 0.0716 | 0.1934 | 0.0811 |
|  | typeS | 85 | 111 | 196 | 0.1008 | -0.0384 | 0.0220 | 0.1259 | 0.0035 | 0.0811 |

**Table S1. Comparison of estimated temporal trends between all, reliable, and unreliable empirical time series across type of errors and taxonomic classes.** For each class and reliability metrics –estimation error (error), statistical power (power), false-positive rate (false post.), Type M error, and Type S error– the table reports the number of time series, mean estimated trend, and median estimated trend for (i) all time series (No. all), (ii) time series classified as reliable according to the thresholds defined in this study (No. realiable), and (iii) time series classified as unrealiable (No. unreraliable).

### References

Collen, B., Loh, J., Whitmee, S., McRae, L., Amin, R., & Baillie, J. E. M. (2009). Monitoring change in vertebrate abundance: The living planet index. *Conservation Biology: The Journal of the Society for Conservation Biology*, *23*(2), 317-327. https://doi.org/10.1111/j.1523-1739.2008.01117.x

Gregory, R. D., & Strien, A. van. (2010). Wild Bird Indicators: Using Composite Population Trends of Birds as Measures of Environmental Health. *Ornithological Science*, *9*(1), 3-22. https://doi.org/10.2326/osj.9.3
